# Genetic immunodeficiency and autoimmune disease reveal distinct roles of Hem1 in the WAVE2 and mTORC2 complexes

**DOI:** 10.1101/692004

**Authors:** William A. Comrie, M. Cecilia Poli, Sarah A. Cook, Morgan Similuk, Andrew J. Oler, Aiman J. Faruqi, Douglas B. Kuhns, Sheng Yang, Alexandre F. Carisey, Benjamin Fournier, D. Eric Anderson, Susan Price, Wadih Abou Chahla, Alexander Vargas-Hernandez, Lisa R. Forbes, Emily M. Mace, Tram N. Cao, Zeynep H. Coban-Akdemir, Shalini N. Jhangiani, Donna M. Muzny, Richard A. Gibbs, James R. Lupski, Jordan S. Orange, Geoffrey D.E. Cuvelier, Moza Al Hassani, Nawal AL Kaabi, Zain Al Yafei, Soma Jyonouchi, Nikita Raje, Jason W. Caldwell, Yanping Huang, Janis K. Burkhardt, Sylvain Latour, Baoyu Chen, Gehad ElGhazali, V. Koneti Rao, Ivan K. Chinn, Michael J. Lenardo

## Abstract

Immunodeficiency often coincides with immune hyperresponsiveness such as autoimmunity, lymphoproliferation, or atopy, but the molecular basis of this paradox is typically unknown. We describe four families with immunodeficiency coupled with atopy, lymphoproliferation, cytokine overproduction, hemophagocytic lymphohistocytosis, and autoimmunity. We discovered loss-of-function variants in the gene *NCKAP1L*, encoding the hematopoietic-specific Hem1 protein. Three mutations cause Hem1 protein and WAVE regulatory complex (WRC) loss, thereby disrupting actin polymerization, synapse formation, and immune cell migration. Another mutant, M371V encodes a stable Hem1 protein but abrogates binding of the Arf1 GTPase and identifies Arf1 as a critical Hem1 regulator. All mutations reduce the cortical actin barrier to cytokine release explaining immune hyperresponsiveness. Finally, Hem1 loss blocked mTORC2-dependent AKT phosphorylation, T cell proliferation, and effector cytokine production during T cell activation. Thus, our data show that Hem1 independently governs two key regulatory complexes, the WRC and mTORC2, and how Hem1 loss causes a combined immunodeficiency and immune hyperresponsiveness disease.

**One sentence summary:** Hem1 loss of function mutations cause a congenital immunodysregulatory disease and reveal its role regulating WAVE2 and mTORC2 signaling.

---

Inborn errors of immunity can paradoxically cause life-threatening infections coupled with lymphoproliferation or autoimmunity by genetically altering global cellular regulatory systems.(*1, 2*) The actin cytoskeleton is a global regulator of cell migration, phagocytosis, immune synapse formation, cell division, vesicle release, and cytotoxicity. Other global cell regulators are the mTOR complexes 1 (mTORC1) and 2 (mTORC2) that control cell metabolism and signaling. How these two systems are molecularly coordinated during immune responses is unknown. The WAVE regulatory complex (WRC), an obligate heteropentamer containing isoforms of the Cyfip1/2, Hem1/2, Abi1/2, HSPC300 and WAVE1/2/3 proteins, dynamically regulates F-actin polymerization for cell migration and immune synapse function and generates a static cortical actin network (CAcN) that controls cell deformation and restricts cytoplasmic vesicle release.(*3–6*) Diverse signals, including the small GTPases Rac1 and Arf1, acidic phospholipids, kinases, and cell surface receptors can cooperatively recruit and activate the WRC.(*7–12*) Rac1 has two binding sites on Cyfip1/2, but none on Hem1. The Arf1 docking site in the WRC is unknown. Genetic diseases have been described for some actin regulatory proteins, including WASP, ARPC1B, and Rac2, but not yet for WRC components. Rapamycin-insensitive mTORC2, a protein complex comprised of mTOR, Rictor, mSIN1, mLST8, Protor1/2, and Deptor, phosphorylates and activates AGC kinases (AKT, SGK1, and PKC) to promote cell survival/proliferation, T cell differentiation, and is required for regulated cell migration through regulation of the actin cytoskeleton.(*13–17*) Thus, the upstream coordination of mTORC2 and WRC-mediated actin polymerization following T cell receptor (TCR) stimulation is important for well-regulated immune response, but still poorly defined.

We investigated five patients, 2 to 16 years old, from four unrelated kindreds, who presented with severe immunodeficiency comprising recurrent bacterial and viral skin infections, septic arthritis, bacteremia, otitis media, and upper respiratory tract infections leading to bronchiectasis (Table S1, Fig. 1A and B left panels, and Fig. S1A, supplemental patient narratives). Patient T cells had selective defects in TCR-induced activation markers CD69 and CD25, proliferation, and IL-2 and tumor necrosis factor (TNF) secretion, though we found surprisingly normal or slightly elevated interferon (IFN)-γ, IL-10, granzyme (Gzm) A/B and perforin production, and CD8 cytotoxicity (Figs. 1C-F, S2H, and S3). Patient cells had reduced Lymphocyte Function-Associated Antigen 1 (LFA-1) inside-out activation, but intact adhesion to immobilized ICAM-1 (Fig. S2A-C). ICAM-1 addition during stimulation improved T cell activation but failed to restore proliferation; additionally, strong TCR stimulation (CD3/28-coated beads), second messenger mimetics phorbol myristate acetate (PMA) and ionomycin (I), or exogenous IL-2 did not rescue activation responses (Fig. S2D-G). Hence, patient T cells have selective defects in response to TCR signaling (Fig. S2I-K). Conversely, the patients also manifested immune system hyperreactivity, which included atopic and allergic disease; increased antibody production (IgM, IgG, and IgE); secondary lymphoid expansion with chronic hepatosplenomegaly and lymphadenopathy (despite normal FAS and TCR-mediated apoptosis), autoantibodies, immune complex glomerulonephritis; and Epstein-Barr virus driven hemophagocytic lymphohistiocytosis (HLH) (Table S1 and S2, Fig. 1B right panel, Fig. S1A-C). Patient T cells showed exaggerated responses to IL-2 stimulation with increased proliferation and secretion by CD4^+^ T cells and elevated perforin and GzmA/B secretion by CD8^+^ T cells (Fig. 1G-I). Immunophenotyping revealed increased B and memory T cells, decreased natural killer (NK) cells, and variable antibody anomalies (poor specific responses but hyper IgG/IgA in two patients and deficient Ig in one patient) (Tables S1, S2 and Fig. S4A).

**Figure 1.**
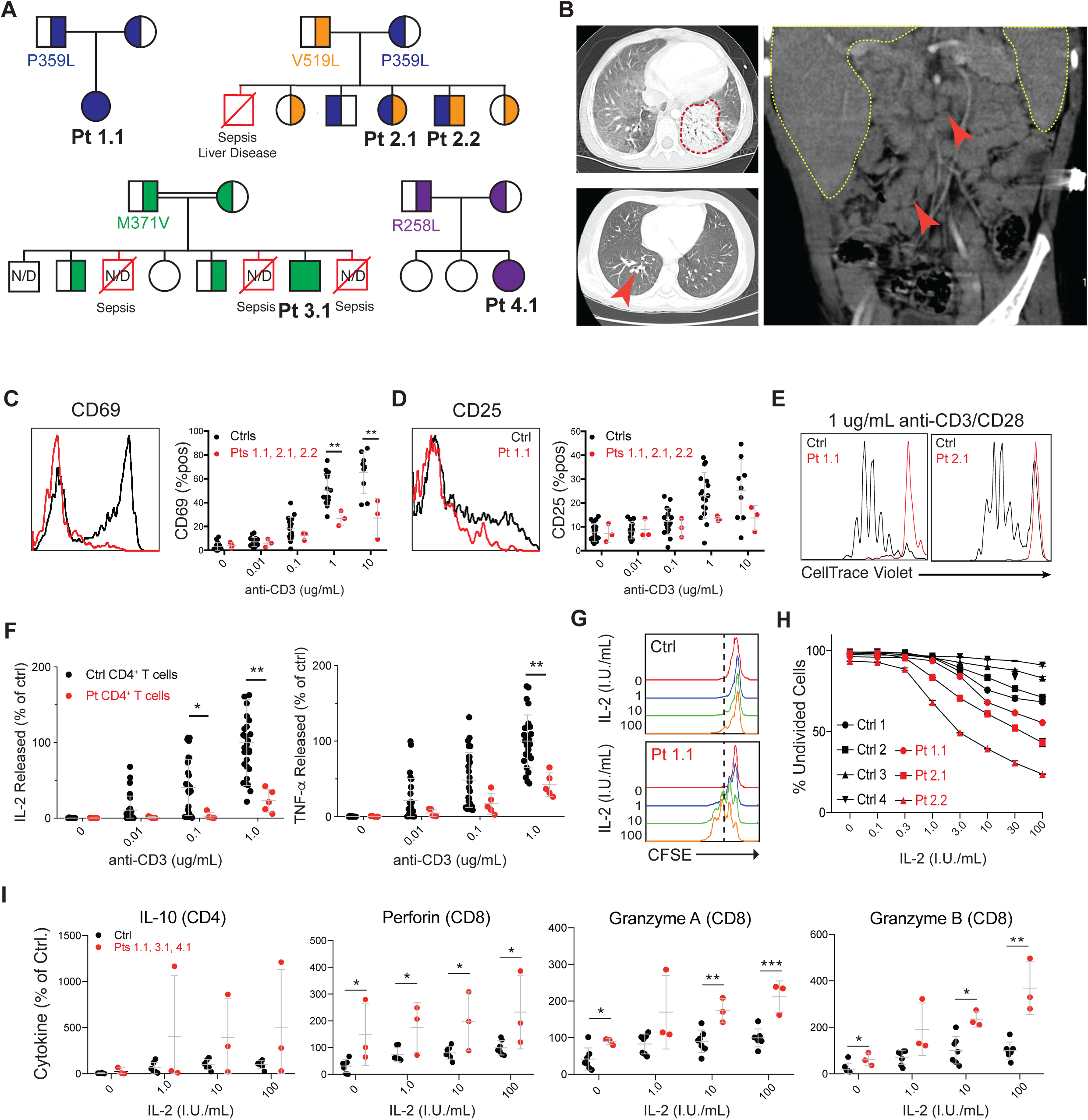
A novel primary immunodeficiency presenting with lymphoproliferative disease, atopic disease, and HLH-like immune activation. **(A)** Patient pedigrees showing recessive inheritance of *NCKAP1L* missense mutations. Deceased siblings of unknown genotype with a similar clinical phenotype are represented by red symbols with their assumed cause of death. **(B)** CT scans of patients demonstrating ground glass opacity and pneumonia (red outline) in Pt 1.1 (upper left), bronchiectasis (red arrow) in Pt. 2.2 (bottom left), and hepatosplenomegaly (yellow outlines) and lymphadenopathy (red arrows). **(C)** CD69 and **(D)** CD25 upregulation on naïve CD4^+^ T cells from patients and controls following stimulation with anti-CD28 and the indicated dose of immobilized anti-CD3. Each dot represents a single control or patient. **(E)** CellTrace Violet plots of naïve CD4^+^ T cells stimulated for 5 days on immobilized anti-CD3/CD28 (1 ug/mL each). **(F)** IL-2 and TNF-a secreted by CD4^+^ T cell blasts upon restimulation for 36 hours with immobilized ICAM-1/anti-CD28 and the indicated dose of anti-CD3. **(G)** T cell proliferation measured by carboxyfluorescein succinimidyl ester (CFSE) dilution of patient CD4^+^ T cell blasts after rest and restimulation with the indicated dose of IL-2 for 96 hours. **(H)** Percent of cells remaining in the undivided peak following restimulation as in **(G)**. **(I)** Soluble cytokine release from patient T cell blasts following 18-hour stimulation with the indicated amount of IL-2. \**P* ≤ 0.05, \*\**P* ≤ 0.01, \*\*\**P* ≤ 0.001, \*\*\*\**P* ≤ 0.0001.

All five patients harbored bi-allelic missense variants in the gene *NCKAP1L,* which encodes Hem1, the only hematopoietic-specific member of the WRC, as the only shared gene defect (Fig. 1A, Fig. 2A, Table S3). The probability of Loss of Function Intolerance (pLI) score for NCKAP1L is 0.99 in ExAC indicating Hem1 is intolerant of sequence changes.(*18*) Hem1 knockout (KO) mice have multiple hematopoietic lineage defects.(*19*) Patient (Pt) 1.1 is homozygous for a missense variant (NM_005337.4:c.1076C>T; p.Pro359Leu). Pt 2.1 and Pt 2.2 are compound heterozygous for P359L along with a second variant (NM_005337.4:c.1555G>C; p.Val519Leu). Pt 3.1 is homozygous for a single nucleotide change (NM_005337.4:c.1111A>G; p.Met37Val). Finally, Pt 4.1 is homozygous for a fourth variant (NM_005337.4:c.773G>T; p.Arg258Leu) (Fig. 1A, Table S3,). These variants were not found in the homozygous state in ExAC, gnomAD or internal databases, are bioinformatically predicted to be deleterious, and the WT amino acids are highly conserved in both Hem1 and Hem2 throughout phylogeny and cluster near the WRC distal Rac1-binding site on, located on Cyfip1 (Fig. S5A, B, C, D).

**Figure 2.**
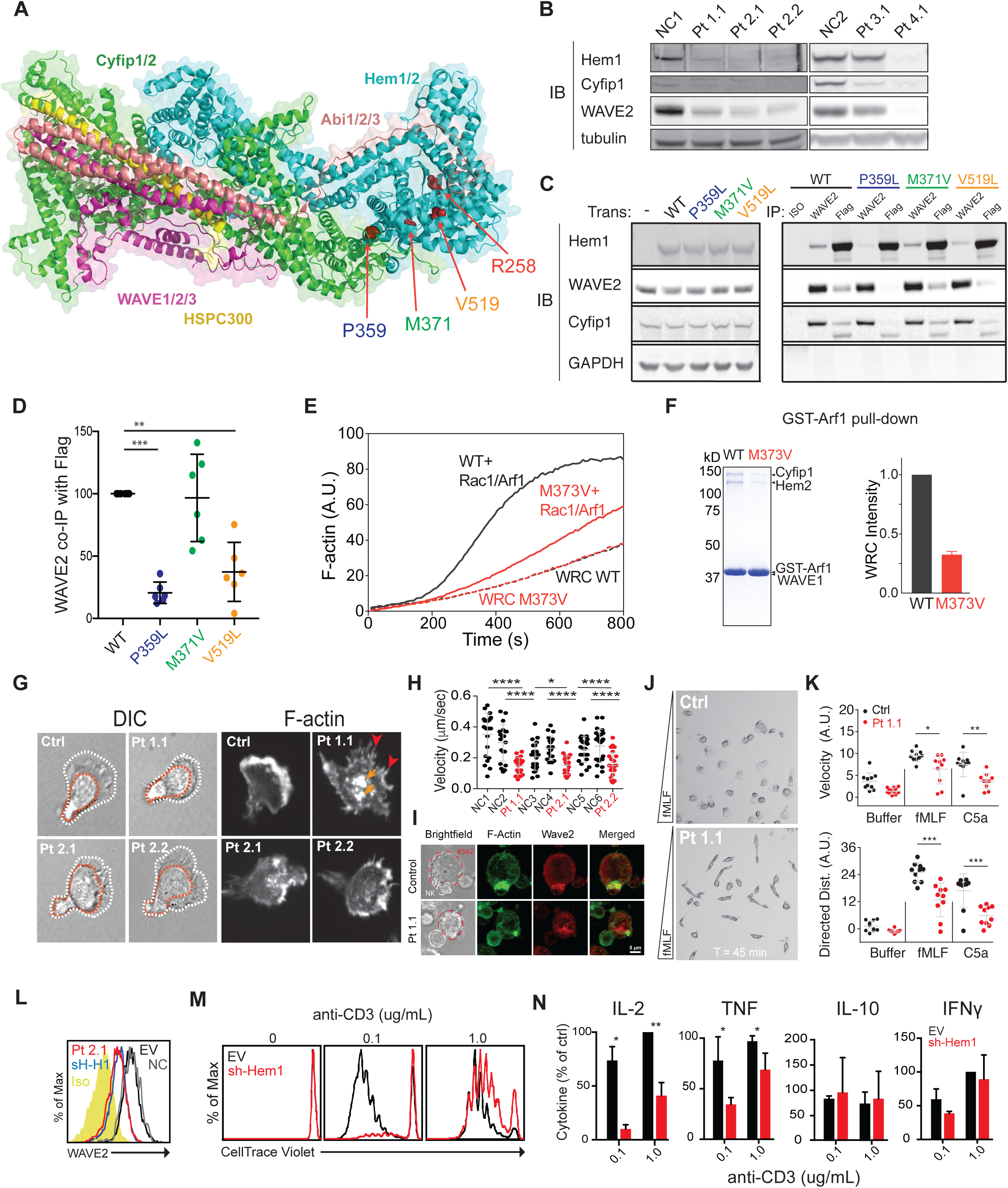
Bi-allelic variants in *NCKAP1L* disrupt WRC stability/function and lead to defective lamellipodia formation and T cell activation. **(A)** Location of patient mutations in Hem1 relative to the overall structure of the WRC (PDB 3P8C)(*10*) (**B**) Immunoblot of WRC components in lysates derived from patient CD8^+^ T cell blasts. (**C**) Representative immunoblot of whole-cell lysates of 293T cells overexpressing the indicated Hem1 construct (left) or immunoprecipitations of the indicated target stained for WRC components (right). (**D**) Quantification of WAVE2 coimmunoprecipitated by Flag-tagged Hem1 constructs as shown in (C). (**E**) Pyrene-actin polymerization assay with WRC230VCA containing Hem2 WT or M373V, with or without activation by a Rac1-Arf1 heterodimer pre-loaded with GMPPNP (See Methods for details). (**F**) Representative Coomassie blue-stained SDS PAGE gel and quantification from 4 repetitions showing GST-Arf1 pull-down of WRC230VCA containing Hem2 WT or M373V in the presence of Rac1 (Q61L/P29S). (**G**) Photomicrographs of CD4^+^ T cell blasts from normal controls (NC) or patients (Pt) spontaneously migrating on ICAM-1 and imaged in real time (DIC) or fixed and stained with phalloidin (F-actin) (right). Red arrows: abnormal formin-bundled actin. Orange arrows: Wiskott-Aldrich syndrome protein (WASp)-mediated actin puncta. (**H**) Spontaneous velocity of individual cells migrating as in (G). (**I**) Representative imaging of immunological synapses between target cells (K562) and NK cells from a normal control or Pt. 1.1, stained for F-actin and Wave2. (**J**) Single frame from Movie S4 showing control and patient neutrophils migrating in a gradient of the chemotactic peptide, N-formyl-L-methionyl-L-leucyl-L-phenylalanine (fMLF). (**K**) Displacement velocity (top) and net directed distance (bottom) in arbitrary units (AU) of 10 randomly selected control or Pt 1.1 neutrophils migrating in chemoattractant gradients (fMLF or C5a) or buffer only. (**L**) Flow cytometry histogram showing Wave2 levels in CD4^+^ T cells from a normal control (NC2) or Pt. 2.2, or normal CD4^+^ T cells transduced with empty vector (EV) or Hem1-shRNA (sh-H1). (**M**) CellTrace Violet dilution histograms showing proliferation of CD4^+^ T cells transduced with EV or shHem1 stimulated on immobilized ICAM-1/anti-CD28 and the indicated dose of anti-CD3. (**N**) Cytokine production by EV or sh-Hem1 transduced CD4^+^ T cells. \**P* ≤ 0.05, \*\**P* ≤ 0.01, \*\*\**P* ≤ 0.001, \*\*\*\**P* ≤ 0.0001.

The R258L, P359L, or V519L mutations caused a loss of Hem1 protein with concomitant loss of Cyfip1 and WAVE2, indicating a destabilized WRC (Fig. 2B and Fig. S6A-C). We confirmed this by making a CRISPR/Cas9 knockout (KO) of Hem1 in Jurkat T cells in which we showed the patient Hem1 mutations (P359L and V519L) recapitulated the WRC loss (Fig. S6D). Moreover, immunoprecipitation of Flag-tagged Hem1 from 293T cells showed that the P359L and V519L mutants had reduced affinity for WAVE2 (Fig. 2C,D). With a cycloheximide translational block, we observed equivalent WT and mutant Hem1 and Wave2 protein turnover suggesting the mutations affected WRC assembly but not intrinsic protein stability (Fig. S6E). The M371V mutation behaved differently, with normal Hem1 and WRC expression in either patient cells or when expressed in Hem1 KO Jurkat cells and immunoprecipitated normally with endogenous WAVE2 in 293T cells (Fig. 2B-D and S6D). We therefore hypothesized that the M371V mutation prevented the binding of a crucial regulator of Hem1/WRC. To test this, we reconstituted the WRC in vitro with recombinant proteins (containing Hem2 with the equivalent M373V substitution, instead of Hem1 which resisted purification) to test the effect of the substitution on ligand binding and activation (Fig. 2E,F and S6F,G).(*3*) We found that Hem2-M373V mutation gave normal yields and chromatographic behavior of the WRC during purification and did not affect WRC binding to the Rac1 GTPase nor its activation by Rac1 (Fig. S6F, G). By contrast, the mutation substantially reduced WRC binding and activation by the Arf1 GTPase in the presence of its cooperative activator Rac1 (Fig. 2E,F and S6F,G). Thus, Hem1/2, specifically the M371/3 residue, is crucial for Arf1 binding and activation of the WRC to promote actin polymerization. Moreover, we used shRNA to deplete Hem1 from healthy donor CD4^+^ T cells and observed the features of patient T cells including WRC loss, defective T cell proliferation and IL-2 / TNF secretion with normal IL-10/IFNγ production (Fig. 2L,M,N). Taken together, we have identified two mechanisms of Hem1 deficiency; R258L, P359L, and V519L act by destabilizing the WRC while M371V leaves the complex intact but disrupts Arf1-WRC binding and activation (Fig. S5E).

Since the WRC generates branched actin filaments underlying the CAcN, lamellipodia, immune synapses (IS), and cell migration, we investigated patient T cells for these functions. (*20, 21*) We observed that patient T cells migrated poorly on ICAM-1-coated surfaces. They exhibited reduced migratory velocities, a loss of lamellipodia and F-actin at the leading edge, and formation of aberrant membrane spikes and F-actin puncta that likely reflect compensatory formin and/or WASp-mediated actin polymerization, respectively (Fig. 2G,H and Movie S1).(*22*) Migrating patient T cells had excessive blebs at their leading edge, which has been previously associated in WRC-deficient cells with lower cortical rigidity (Movie S1).(*23*) Live cell imaging of T cell immunological synapses formed by either patient cells or Hem1 KO Jurkat T cells reconstituted with mutant Hem1 alleles also showed abnormal lamellipodia, increased formin and WASp-dependent actin structures, and diminished synapse stability (Movies S2 and S3). Defective F-actin accumulation was also observed in the patient NK cell-target synapse, though cytolytic activity was largely preserved, possibly due to WASp-dependnet actin polymerization (Fig. 2I and S4B-D). (*24, 25*) Like Hem1-KO mouse neutrophils, dendritic cells or HL-60 cells, we found that patient neutrophils also migrated poorly in chemoattractant gradients and exhibited actin defects (Fig. 2J,K, and S7A-D, Movie S4).(*19, 21, 26*) Strikingly, we observed that patient neutrophils could not form a broad leading edge or maintain directional persistence and became dramatically elongated with competing leading edges moving parallel to or against the chemotactic gradient (Movie S4). Defective neutrophil migration likely contributes to the bacteremia and recurrent skin infections in our patients.

We next investigated how Hem1 deficiency caused immune hyperactivation. Both patient cells and Hem1-KD T cells exhibited high constitutive and IL-2 induced GzmA/B secretion and patient B cell lymphoblasts (BLCLs) secreted more IL-10 than controls (Fig. 3A and S8). However, IL-2 receptor gamma chain trafficking or STAT phosphorylation were unaffected in Hem1-deficent cells (Fig. S9).(*27*) Since the CAcN can potently restrain lytic granule membrane fusion, degranulation, and exocytosis of Lamp1/CD107a, we examined whether Hem1-deficiency caused increased degranulation. (*28*) Indeed, we found that patient T cells expressed more surface CD107a than control cells, both at baseline and after IL-2 or PMA/I stimulation. (Fig. 3B) We also observed a 25% decrease in average cell spreading area and F-actin polymerization at immune synapses formed by patient T cells, showing that the CAcN network is thinner and may be more easily traversed in Hem1-deficient cells (Fig. 3C,D). We verified this conjecture by gently thinning the CAcN using the actin-depolymerizing agent latrunculin A (LatA) and found CD107a surface expression was augmented in cells stimulated with IL-2 and dramatically increased with PMA/I stimulation (Fig. 3E, top). Interestingly, LatA concentrations that increase CD107a and GzmA/B exocytosis (10 nM) were over 10-fold below those required to prevent cell spreading or T cell activation (0.1-1uM) (Fig. 3E, bottom and 3F). Thus, we conclude that Hem1/WRC regulates the CAcN to control degranulation and cytokine release by T cells responding to antigen, and possibly by bystander T cells in response to IL-2. In patients, loss of CAcN integrity likely causes the patients’ autoimmune, lymphoproliferative, and HLH-like phenotypes, at least in part through increased degranulation and cytokine release. Given that IL-2 can cause hyper-cytokine release, we tested Jak inhibitors and found they prevent this responses in the Hem1-KD cells (data not shown).

**Figure 3.**
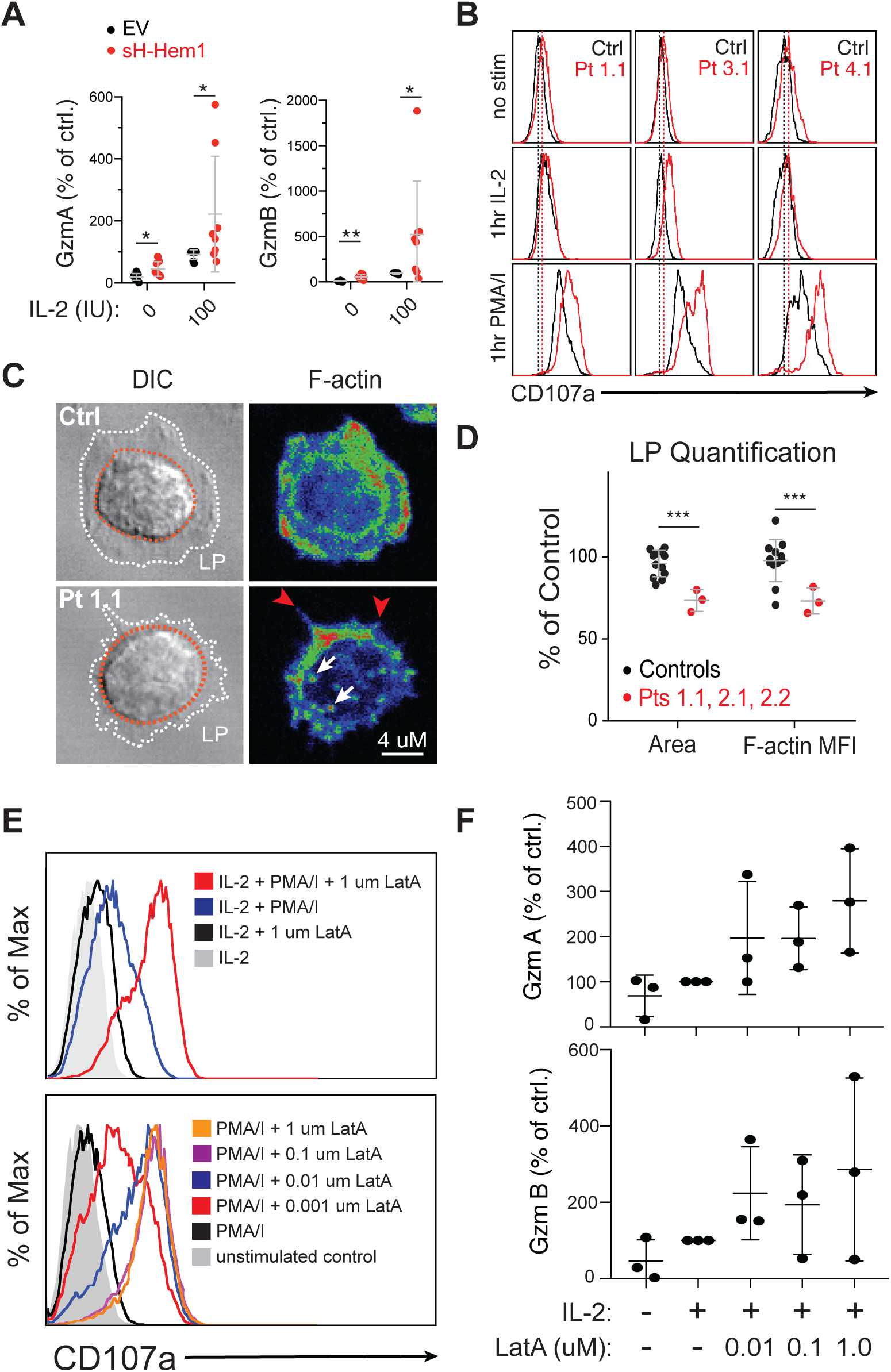
Hem1 deficiency and reduced F-actin leads to IL-2 induced cytokine release and increased mitogen-induced degranulation. (**A**) Soluble granzyme A (GzmA) and B (GzmB) release from normal control T cell blasts transduced with either empty vector (EV) or shRNA against Hem1 (shHem1) following 18-hour stimulation with the indicated amount of IL-2. Each dot represents a different donor. (**B**) Surface CD107a following 1-hour stimulation under the indicated conditions in patient and control T cell blasts. (**C**) Representative photomicrographs of a CD4^+^ T cell blast from either a normal control (NC) or Pt 1.1 spreading on stimulatory coverslips stained with phalloidin. The lamellipodial (LP) region is highlighted between the white and red rings in the differential interference contrast (DIC) microscopy image. (**D**) Quantification of total cellular area and F-actin mean fluorescence intensity (MFI) for the average of patient and control cells over multiple experiments. Each dot represents the cell average for an individual patient or control. (**E**) Surface CD107a following 1-hour stimulation as indicated in the presence or absence of 1 μM latrunculin A (LatA) (top) and surface CD107a following 1-hour PMA/I stimulation and simultaneous treatment with the indicated dose of LatA (bottom). (**F**) Soluble granzymes released from control T cell blasts treated with 100 I.U./mL IL-2 and the indicated amount of LatA for 18 hours. \**P* ≤ 0.05, \*\**P* ≤ 0.01, \*\*\**P* ≤ 0.001

To further understand how Hem1 abnormalities caused immunodeficiency, we studied TCR-induced signaling events. We found that Hem1 mutations caused no defect in proximal TCR signaling events including TCR-stimulated Zap70, PLCγ-1, ERK1/2 phosphorylation and TCR- or thapsigargin-induced Ca^2+^ flux, similar to reports of Wave2-deficient T cells (Fig. 4A,B and Fig. S10).(*20*) However, both patient and Hem1-KD T cells had a specific defect in the phosphorylation of AKT at Ser473 after TCR stimulation (Fig. 4C, D and Fig. S11A-D). AKT Ser473 is a well-known substrate of the mTORC2 complex.(*29*) We confirmed this by adding an mTOR catalytic inhibitor (Ku 0063794) during TCR stimulation which blocked AKT Ser473 phosphorylation and other mTOR-dependent phosphorylation events (Fig. 4E and Fig. S11E). By contrast, LatA, which reduces the CAcN, had the opposite effect: it enhanced the phosphorylation of AKT, ERK, and ribosomal protein S6, indicating that the reduced AKT Ser473 phosphorylation in Hem1-deficient cells was independent of Hem1 function in the WRC. To determine if Hem1 or WAVE2 interacted with proteins involved in the phosphoinositide 3-kinase (PI3K)/AKT pathway, we performed a series of immunoprecipitation (IP)-mass spectrometry experiments to compare the interactomes of Wave2, Hem1-WT, and Hem1-M371V in Jurkat cells and control CD4^+^ T cell blasts (Fig. S12A, Table S4). We readily detected all known members of the WRC among 41 identified WRC interacting proteins common to Jurkat cells and primary T cells using both Hem1-Flag IP and WAVE2 IP. We did not observe a significant difference in interactomes between the M371V and WT-Hem1 (Fig. S12B-E). We also detected several mTORC2 components, including mTOR and Rictor in the Hem1 IP but not in control or WAVE2 IPs (Table S4 and Fig. 4F). This raised the possibility that Hem1 interacted with mTORC2 outside of the WRC. We therefore immunoprecipitated Hem1-Flag (WT/P359L/M371V), Flag-GFP, myc-Rictor, or WAVE2 that were expressed in 293T cells and observed that Hem1 and Rictor specifically co-immunoprecipitated with each other, however WAVE2 and Rictor did not co-precipitate (Fig. 4G, S13A). Interestingly, the P359L Hem1 coimmunoprecipitated Rictor better than the WT-Hem1, suggesting that disrupting the Hem1-WAVE2 interaction might liberate the mutant Hem1 to better interact with Rictor. Thus, Hem1 interacts with Rictor independently of the WRC and is required for mTORC2 activity downstream of the TCR. Importantly, Hem1-deficient patient cells expressed normal levels of Rictor, suggesting that Hem1 is not required for the stability of Rictor or mTORC2 (Fig. S13B). To determine if inhibition of mTORC2 could recapitulate the patient cellular phenotype, we targeted Rictor by shRNA and observed that T cell proliferation and IL-2 and TNF secretion were markedly impaired (Fig. 4H,I and Fig. S13C). Additionally, inhibitors of the PI3K, AKT, or mTOR kinases, but not low dose LatA impaired T cell proliferation and IL2 and TNF secretion (Fig. 4J,K and Fig. S13E,F). Critically, rapamycin treatment, which specifically targets mTORC1, had less effect than the mTOR catalytic inhibitor, suggesting that mTORC2, but not mTORC1, was obligatory for these T cell responses. Moreover, we found that PI3K/AKT/mTOR inhibition had little effect on CD69 upregulation or secretion of perforin and GzmA/B, essentially phenocopying the defects observed in patient cells (Fig. S13D,F). Together, our data reveal a new role for Hem1 as an upstream regulator of mTORC2 catalytic activity, independent of its role in the WRC (Fig. 4L).

**Figure 4.**
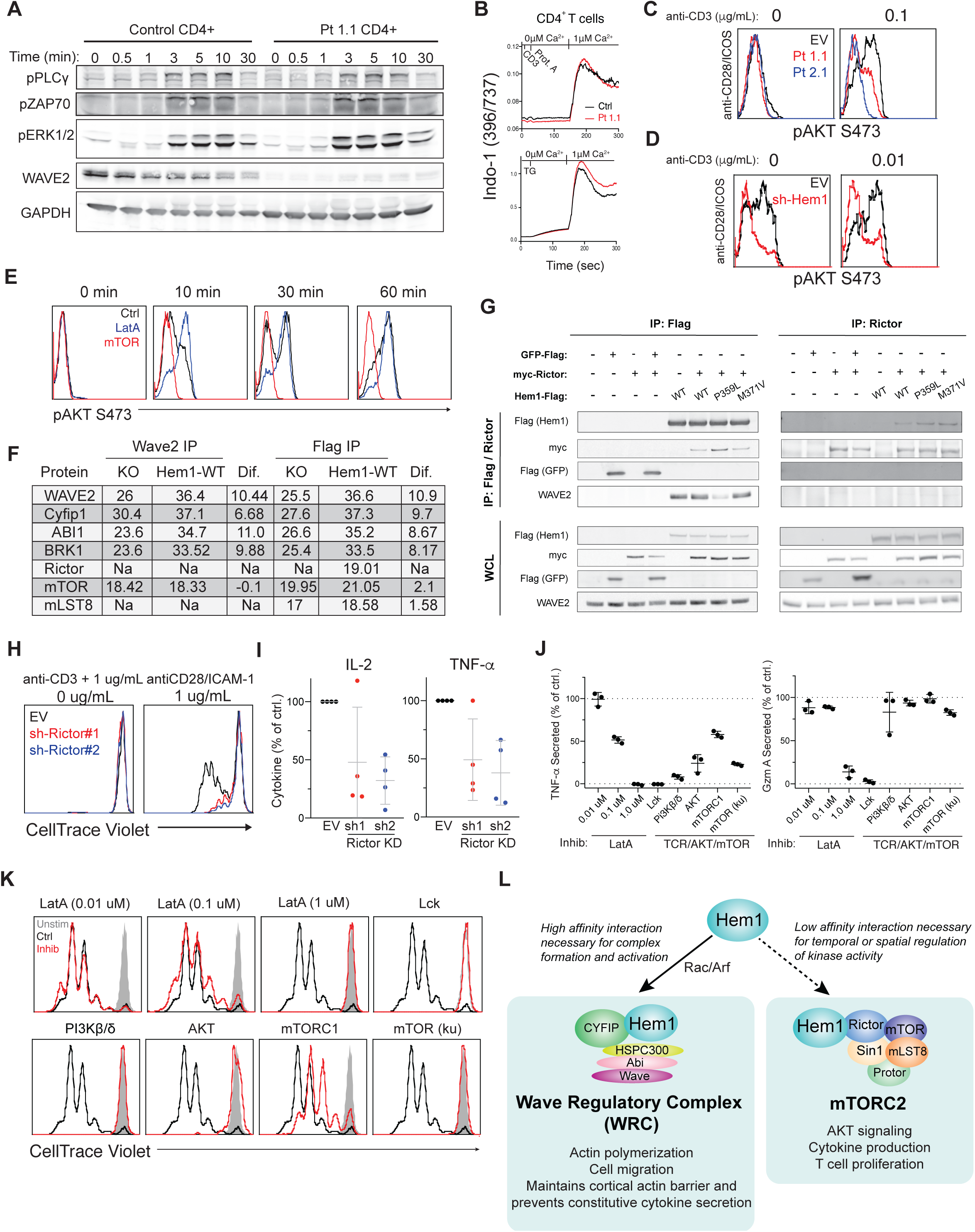
Hem1 is required for proper mTORC2 functionality and downstream T cell effector functions. (**A**) Immunoblot of lysates from normal control and patient T cells restimulated for the indicated times with soluble anti-CD3/CD28. (**B**) Calcium flux in patient and normal control T cells following crosslinking with anti-CD3/28 and Protein A (Prot. A) or thapsigargin (TG)-induced ER store release. (**C**) Representative flow staining for AKT phosphorylated on Ser473 (pAKT S473) in patient or control CD4^+^ T cells stimulated for 10 minutes with anti-CD28/ICOS (1 μg/mL each), and the indicated dose of anti-CD3. (**D**) Representative flow plots showing pAKT S473 in CD4^+^ T cells from healthy donors transduced with either EV or sh-Hem1 and stimulated as in (C). (**E**) Representative flow plots of AKT S473 in CD4^+^ T cells treated with either 1 μM LatA or 10 μM Ku-0063794 (mTOR) prior to stimulation with anti-CD3/CD28/ICOS for the indicated time. (**F**) WRC and mTORC2 components identified by IP-MS analysis. Values indicate log_2_(intensity). (**G**) Flag and Rictor IP from 293T cells transduced with myc-Rictor and either Flag-tagged GFP or Flag-tagged Hem1 (WT or mutant) and blotted for the indicated targets. (**H**) CellTrace violet plots showing proliferation of naïve CD4^+^ T cells transduced with empty vector (EV) or shRNA directed against Rictor (sh-Rictor#1 and #2). (**I**) Cytokine secretion by control and Rictor knockdown (KD) CD4^+^ T cells following 18-hour restimulation. (**J**) IL-2 and Gzm A secretion by CD8^+^ T cell blasts restimulated as in (I) in the presence of the indicated inhibitor. (**K**) Proliferation of normal control T cells treated with the indicated inhibitors and stimulated as in (H). (**L**) Model illustrating the role of Hem1 in regulation of WRC- and mTORC2-mediated cellular functions.

Our data show that LOF mutations in *NCKAP1L*, encoding Hem1, disrupt WRC-mediated actin polymerization and abrogate mTORC2 activation of AKT. This results in an autosomal recessive inborn error of immunity affecting multiple hematopoietic lineages and causing susceptibility to bacterial/viral infections, atopic disease, autoimmunity, cytokine overproduction and lymphoproliferation. The WRC plays a vital role in building branched actin networks and the CAcN by stimulating the actin-nucleating activity of the Arp2/3 complex downstream of Rac1 and Arf1. Hem1 is the sole lymphoid-specific component of the WRC, but how it promotes WRC functions has not been previously described. We now show that Hem1 is required for Arf1 binding and Arf1-dependent WRC activation which we observed were specifically impaired by the M371V amino acid substitution. We also show that Hem1 and the WRC are essential for the CAcN, which governs cytokine secretion and lytic granule fusion with the plasma membrane. Hem1-deficient patients have prominent immune hyperactivation, which we can now attribute, at least in part, to increased vesicle release. Finally, we uncovered a critical interaction between Hem1 and Rictor that regulates mTORC2 activity in response to TCR stimulation. The Hem1-Rictor interaction appears to have previously escaped detection possibly because it is a low affinity interaction (Fig. 4F, G) and because the commonly used 293T cells do not express the hematopoietically-restricted Hem1.(*30*) Loss of Hem1 or Rictor, or pharmacological inhibition of mTORC2, markedly impaired AKT S473 phosphorylation and specific TCR-induced responses. We posit that Hem1 coordinates WRC-mediated actin polymerization and mTORC2 catalytic activity in response to signals that activate both protein complexes, such as PI3K and Rac1, during TCR signaling. This mechanism sheds new light on previous data showing that mTORC2 can be activated downstream of actin-generated membrane tension and may provide negative feedback regulation of the WRC.(*31*) Thus, this human immunodysregulatory disorder reveals new and unexpected roles for the highly conserved actin regulator Hem1, which may lead to the development of new valuable immunotherapies.

## ACKNOWLEDGMENTS

The authors would like to thank Andrew Oler, Ke Huang, Sandhya Xirasagar, Darrell Hurt, and other members of the Bioinformatics and Computational Biosciences Branch (BCBB), NIAID for variant assessment and bioinformatics support. Finally, we thank the patients and family members for participating in this study, and without whom it would not be possible to conduct this research. We thank Helen Su for scientific input and discussions and for careful reading of the manuscript.

## AUTHOR CONTRIBUTIONS

WAC, SAC, and AJF performed experiments related to WRC expression/function, T cell activation/function, analyzed data, and interpreted results. SAC performed experiments related to the functional validation of P359L, M371V, and V519L. MCP, AVC, AFC, EMM, and JSO directed or performed NK cell experiments, analyzed data, and interpreted results. DBK performed neutrophil experiments and analyzed data. WAC prepared IP-Mass Spec samples and DEA performed Mass-Spec analysis and generated the list of interacting proteins. SY performed in-vitro WRC reconstitution and actin polymerization assays. SP, GC, and VKR oversaw care of Pt 1.1 and VKR, MS, and AO performed and interpreted WES for Kindred 1. JWC, and NR oversaw care of Pts 2.1 and 2.2 and TNC, ZHC-A, SNJ, DMM, RAG, and JRL performed and interpreted WES to identify causal variants for kindred 2. MAH, NAK, ZAY, SJ, and GE oversaw care of Pt 3.1 and GE performed and GE and AO interpreted WES for kindred 3 to identify causal mutations. WAC(2), BF, and SL oversaw care of Pt 4.1 and performed and interpreted WES to identify causal mutations. Patient clinical histories were prepared by WAC, MCP, and attending physicians. JSO, LRF, JKB, SL, BC, GE, VKR, IKC, and MJL supervised various aspects of the project and project personnel. WAC, MCP, SAC, IKC, and MJL interpreted results and wrote the manuscript. WAC took day-to-day responsibility for the study. MJL coordinated the overall direction of the study. All authors read and provided appropriate feedback on the submitted manuscript.

## Materials and Methods

### Methods

#### Study participants and human sample collection

We enrolled five patients from four families from living in Canada, the United States, the United Arab Emirates, and France. Patients were included in the study based on a molecular identification of function-affecting variants in *NCKAP1L*, irrespective of age and sex. The ages of the patients ranged from 2 to 17 years old as of June 2019. Patients were initially identified on the basis of recurrent infections, EBV-driven HLH, and lymphoproliferative disease. They underwent whole exome sequencing following initial clinical evaluation and in three cases, negative screening results for ALPS-causal mutations. For details regarding individual patients, see the individual patient narratives in the supplemental material available with the full text of this article. All human subjects (or their legal guardians) provided written informed consent in accordance with Helsinki principles for enrollment in research protocols that were approved by the Institutional Review Boards of the National Institute of Allergy and Infectious Diseases, National Institutes of Health (NIH), Baylor College of Medicine, and collaborating institutions. Patient samples were obtained from the NIH Clinical Center or other institutions overseeing patient care under approved protocols and shipped to the NIH. Healthy control blood or buffy coats were obtained at the NIH Clinical Center under approved protocols. Mutations will be archived by Online Mendelian Inheritance in Man (OMIM) at time of publication, and whole-exome data will be or already have been submitted in dbGaP.

#### Genomic analysis

Genomic DNA (gDNA) was obtained from probands and family members by isolation and purification from peripheral blood mononuclear cells (PBMCs) using Qiagen’s DNeasy Blood and Tissue Kit or similar kit. DNA was then submitted for Whole Exome Sequencing (WES) through either the NIH Clinical Center, BCM, or via a commercial service. WES was performed on all patients and was combined with homozygosity/ loss of heterozygosity mapping. All sequenced DNA reads were mapped to the human genome reference. Single nucleotide variant and indel calling were performed. All SNVs/indels were annotated, filtered, and prioritized for autosomal recessive putative disease-causing variants based on the pedigree analysis.

#### Cells, Media, and Cell Culture

Stimulation Buffer (Hepes buffered saline, 1% BSA, 0.1% FBS, 2mM MgCl_2_, 1mM CaCl_2_, 6mM Glucose) was used for all in-vitro short term stimulations. DMEM, RPMI, or X-Vivo15 media were supplemented with 10% Fetal Bovine Serum, 1% L-Glutamine (2mM), 1% Penn/Strep (all Invitrogen) and 50 μM 2-ME (Sigma) to make complete media. Complete RPMI was supplemented with 100 I.U./mL IL-2 (Rouche) for primary cell culture or 20% FBS for growth of BLCL lines. HEK293T cells and Jurkat E6.1 cells were from ATCC and were cultured in complete DMEM and complete RPMI, respectively. LentiX-293T were purchased from Clontech and were cultured in complete DMEM. The mouse mastocytoma P815 target cell line was from ATCC and cultured in complete DMEM. Lymphoblastoid cell lines were generated from patient PBMC by transformation with B95-8 EBV cell supernatant and cultured in RPMI medium with 20% FBS, 2 mM glutamine, and 1 μg/ml cyclosporin A. K562 cell lines used as target cells for Cr51 release assays and NK cell conjugate analysis by confocal microscopy were maintained in RPMI media supplemented with 10% FCS. Peripheral blood mononuclear cells were isolated from anti-coagulated blood by ficol paque density separation by centrifugation (800xg/20minutes) followed by collection of the buffy coat layer and extensive washing in PBS. PBMCs were then immediately used or frozen at 20-50×10∧6/mL in freezing media (90%FBS and 10% DMSO). Human T cells were isolated from fresh or frozen PBMCs using the appropriate kit from Miltenyi Biosciences according to the manufacturer’s protocol (Naïve CD4^+^, Naïve CD8^+^, PanT, and Pan CD4^+^). Purity of cells was routinely verified by flow cytometry. Human T cells were then either used immediately for functional studies or activated by CD3/28 magnetic beads, 1 bead/ 2 cells, (Invitrogen) and expanded in complete RPMI supplemented with IL2. T cell blasts were used for functional analysis within 3 weeks following activation. Patient and control neutrophils were isolated by the neutrophil monitoring lab (NIH) by density gradient separation and used immediately for experiments.

Hem1 or Wave2 deficient Jurkat E6.1 Cells were generated by CRISPR RNP-mediated deletion followed by single cell cloning. Guide RNAs against Hem1 and Wave2 were generated by using IDT’s online tool and the top two sequences were pooled following resuspension for a 100uM final stock solution. Guide RNA was mixed 1:1 with fluorescent tracrRNA (IDT) and heated to 95°C for 5 minutes then allowed to cool to room temperature. 9 μL of the tracr/guide complexes were added to 3 μL of purified Cas9 protein (30 μg) and 3 μL of duplex buffer (both IDT). 10×10^6^ Jurkat T cells were electroporated using the P2 primary Cell 4D-nucleofector kit from Lonza with program EH-100. Cells were allowed to recover, and then single cell cloned by serial dilution and screened for loss of Hem1 or Wave2. Cell lines were confirmed as mycoplasma negative before their use in experiments using a PCR-based assay (Sigma Aldrich).

#### Plasmids and Cloning

The pLV-EF1a-IRES-puro, GFP-Flag expression vector, Myc-Rictor expression vector, pcDNA3.1-ccdB-3xFLAG-V5, and the two Rictor shRNA vectors were purchased from Addgene. The pEnter/D-TOPO entry vector was purchased from Thermo Fischer. shRNA vectors against Hem1, the lentiviral packaging constructs (pMD2.G and psPAX2), and the Lifeact-GFP expression construct were a kind gift from Janis K. Burkhardt.

Full length Hem1 was PCR-amplified from cDNA prepped from T cells from a healthy control using Qiagen’s RNeasy kit and the SuperScript-III First Strand Synthesis System for RT-PCR (Invitrogen) using random hexamers and the manufacturer’s protocols. To amplify full length Hem1, touchdown PCR amplification with Phusion Flash polymerase (Thermo Fisher) was performed using with the following primers, F-5’-C<u>ACCATGT</u>CTTTGACATCTGCTTAC-3’, and reverse, 5’-CAGTTTAGGTGGAAGGCCCGAG-3’. The PCR product was gel purified and directionally cloned into the pENTR/D-TOPO Gateway cloning vector using the the pENTR/D-TOPO Cloning Kit with One-Shot/TOP10 chemically competent cells (Thermo Fisher). The WT-sequence was verified by Sanger sequencing. The P359L, M371V, V519L mutants were produced by site-directed mutagenesis of the WT sequence using the following primers:

P359L:

F-5’-GATGAACTGGGACTACTGGGTCCTAAGGCT-3’

R-5’-GTAGTCCCAGTTCATCAGCCAACACAGTCTC-3’

M371V:

F-5’-TGCTTTCGTGGCCCTGTCCTTCATTCGTG-3’

R-5’-CAGGGCCACGAAAGCAAAAAGAGCCTTAGG-3’

V519L:

F-5’-GGAGTGGAAGAGAATGAGGTTCATCACCTTGGC-3’

R-5’-GCCAAGGTGATGAACCTCATTCTCTTCCACTCC-3’

Hem1 expression constructs containing C-terminal 3xFLAG-v5 tags were generated in a pcDNA3.1 backbone (Addgene) using Gateway cloning technology (Thermo Fisher) as follows. First, WT Hem1 was PCR amplified using the WT forward primer listed above and a reverse primer that removed the stop codon: (R-5’-GTTTAGGTGGAAGGCCCGAGACACCT -3’). This product was gel purified using the NucleoSpin Gel and PCR Cleanup Kit (Machinery-Nagel) and cloned into the pENTR/D-TOPO Gateway cloning vector via the BP-Clonase reaction. Finally, Hem1 sequences were shuttled to the pcDNA3.1 destination vector containing a C-terminal 3xFLAG-v5 tag via the LR-Clonase reaction.

Lentiviral constructs were generated in a pLV-EF1a-IRES-puro backbone using InFusion cloning technology (Takara Bio). Briefly, C-terminally FLAG-tagged Hem1 sequences were amplified using the following primers: (F-5’-TAGATCGCGAACGCGATGTCTTTGACATCTGCTTACCAGC-3’; R-5’-CGCCCTCGAGGAATTTGTAGAATCGAGACCGAGGAGAGG-3’). After gel purification, Hem1 PCR products were incubated with the InFusion Enzyme and linearized pLV-EF1a-IRES-puro backbone previously digested with EcoRI and MluI. All constructs were verified by Sanger sequencing.

#### Plasmid Transfection and Lentiviral Production/Transduction

For Hem1 overexpression experiments, 2×10^6^ HEK293T cells were transiently transfected with 20 μg Hem1-3xFLAG plasmid DNA using Lipofectamine 3000 reagent (Thermo Fisher) and cultured for 72 hours before harvest. For Rictor overexpression experiments, 2×10^6^ HEK293T cells stably expressing FLAG-tagged Hem1 were transiently transfected with 2 μg myc-Rictor plasmid (Addgene). Control HEK293T cells were transfected with myc-Rictor or 10 ng GFP-3xFLAG as described. Lifeact-GFP was transiently transfected into primary T cell blasts using the Amaxa 96-well shuttle system (Lonza) and the recommended protocol for activated human T cells.

For production of lentiviruses, LentiX-293T cells (Takara Bio) were grown to 80% confluency and transfected with 4 μg pMD2.G, 14 μg psPAX2, and 18 μg of lentiviral expression or knockdown vectors (using Lipofectamine 3000 reagent (Invitrogen)), according to the manufacturer’s protocol. Culture supernatants containing lentiviruses were harvested after 72 hours, clarified by centrifugation for 15 minutes at 1500 rpm, and stored at -80°C until use.

For generation of stable Hem1-mutant expressing Jurkat cells, Hem1-knockout Jurkat cells were seeded in 24-well plates in RPMI supplemented with 8 μg/mL Polybrene (Sigma) and 1 mL of Hem1-containing lentiviruses. Plates were centrifuged for 2 hours at 800xg at 32°C. Transduced cells were cultured in complete RPMI for 48 hours before being maintained in 1 μg/mL puromycin selection for at least 48 hours. The same procedure was used to transduce Jurkat cell lines with LifeAct-GFP, HEK293T cells with Hem1-3XFLAG constructs, and primary T cells with shRNA constructs.

#### Inhibitors and Inhibitor Treatments

Inhibitors were added to T cells 30 minutes prior to restimulation and kept in culture during the duration of the T cell stimulation. A vehicle-only control was included equivalent to the highest amount used in the inhibitor-treated cells. The PI3K*β*/*δ* inhibitor AZD 8186 was purchased from Selleckchem and used at a final concentration of 100 uM, The AKT inhibitor MK-2206 H-Cl was purchased from Selleckchem and used at a final concentration of 10 uM. Rapamycin was purchased from Selleckchem and used at a final concentration of 1uM. The mTOR-catalytic inhibitor KU-0063794 was purchased from Selleckchem and used at a final concentration of 10 uM. Latrunculin A was purchased form Sigma and used at the indicated concentrations. Lck inhibitor was purchased from Millipore and used at a final concentration of 10 uM. All inhibitors were resuspended in DMSO and single use stock solutions were kept at -20 C.

#### Neutrophil Studies

Neutrophils were isolated from heparinized blood by standard procedures. Chemotaxis assays were performed as follows: isolated PMNs (5 × 103 cells) and either control, C5a, or, fMLF were added to appropriate wells of an EZ-TAXIScan instrument. Digital images were acquired every 30 seconds for 1 hour. Images were converted to stacks using ImageJ software (version 1.46r, National Institutes of Health), MTrackJ plug-in was used to track individual cell migration, and track measurements were analyzed using Microsoft Excel. Alternatively, isolated PBMNs were added to the top chamber of a transwell migration plate (ChemoTx) with the indicated concentration of the appropriate chemoattractant in the bottom well. Cells were allowed to transmigrate at 37 C, plates were then collected and % migration was determined. For microscopy of neutrophils form patient 1.1, cells were stimulated in solution, cyotospin and fixed to coverglass slides, permeabilized and stained with phalloidin to visualize the F-actin network.

#### CD8^+^ T cell Cytotoxicity

CD8^+^ T cell blasts from normal controls and patients were collected, rested for 3hrs in basal RPMI, and resuspended to 1×10^6^/mL and serial dilutions were prepared for addition of target cells. P815 cells (ATCC) were collected by gentle lifting in PBS-/-with EDTA and a single cell suspension was generated by repeat pipetting and passage through a 70 uM filter. Cells were then either left alone or coated with 1 ug/mL anti-CD3 for 30minutes on ice in basal RPMI. Cells were washed 3X with complete media and resuspended to 1×10^6^/mL in complete RPMI. 5×10∧4 target cells were then added to a 96 well V-bottom plate in triplicate for each dose/ donor. An appropriate number of T cells were then added to reach the appropriate effector to target ration from the serial dilutions generated earlier. Volume/ well was held constant at 200uL. For the no CD3 coated target cells just do the maximal effector to target ratio. A no cell control, effector only spontaneous lysis, a target cell spontaneous lysis only control, and a maximal target cell lysis control were included. Cells were allowed to incubate at 37 C for 4-6hrs. Cytotoxicity was then measured from cell supernatants based on release of lactate dehydrogenase (LDH) released from dying cells based on the CytoTox 96® Non-Radioactive Cytotoxicity assay from Promega and done according to manufacture protocol. Percent cytotoxicity was calculated as described in the product insert and normalized to the results obtained from the normal control cells to average across multiple experiments.

#### ^51^Cr release assays to analyze NK cell-mediated cytotoxicity

Chromium release assays were used to determine NK cell cytotoxicity as previously described.(*32*) Frozen PBMCs were thawed, left to rest for 3 hours at 37°C and 5% CO_2_. Resting cells were counted and incubated at varying ratios (50:1, 25:1, 12.5:1, 6.25:1, and 3.13:1) with 10^4^ K562 target cells that were previously labeled with 50 mCi of ^51^Cr and then washed. Where indicated, assays were performed in the presence of 1000 U/mL IL-2 (Roche). Effector-target conjugates were incubated in 200 mL in round-bottom 96-well plates (Corning, Corning). After 4 hours of incubation, positive controls for maximal lysis were produced by lysing labeled target cells with 1% octylphenoxypolyethoxyethanol. Supernatants were harvested and transferred to LumaPlates (PerkinElmer). The supernatant was dried, and plates were read on a TopCount gamma counter (PerkinElmer). Values were plotted using GraphPad Prism 7 software. Values were normalized to the percentage of NK cells in the initial PBMC population to calculate the lytic units.

#### NK synapse analysis

Conjugate staining was performed as previously described.(*33, 34*) Briefly, PBMCs from patients were incubated with susceptible K562 target cells for 15 minutes in a conical tube at 37°C and 5% CO_2_ and subsequently mounted on a slide for 20 minutes at 37 C and 5% CO_2_ to facilitate conjugate formation. After incubation, cells were fixed and permeabilized and then stained intracellularly with anti-tubulin biotin (Life Technologies), and Wave2 rabbit primary antibody (Cell signaling) followed by streptavidin–Pacific Blue (Life Technologies) and Alexa Fluor-568 goat antirabbit secondary antibody as well as anti-perforin–Alexa Fluor 488 (BioLegend) and phalloidin–Alexa fluor 647 (Life Technologies). Slides were mounted using ProLong Glass antifade reagent (ThermoFisher Scientific) and #1.5 glass cover slips (Corning) and cured for 24 hours prior to observation by confocal microscopy. Fixed cells were imaged using a Zeiss Axio Observer Z.1 microscope stand equipped with a Yokogawa CSU10 spinning disk and a Hamamatsu Orca-R2 C10600 CCD camera. Laser lines used include 405, 488, 561 and 647 nm Coherent OBIS LX powered by a MultiLine LaserBank (Cairn Research). Emitted light were selected using the following filters as appropriate for the relevant dye: 460/50, 520/35 and 593/40 (Chroma Technology Corp). Imaging was performed using the plan apochromatic 63× 1.4NA oil immersion objective and MetaMorph software (v. 7.8.13; Molecular Devices) was used for hardware control and image acquisition. Images were taken in multiple Z-planes and stacked for analysis. Raw 3D stacks of images were exported for processing and analysis in Fiji (*35*) using a custom script. Briefly, binary masks of individual cells were created using the threshold tool applied to the F-actin detection fluorescent channel. Cell outlines were filtered using a size cut-off (>10µm^2^). The fluorescence intensity of the Actin and Wave2 staining channels for each cell was then measured for the sum of all planes of the 3D stack within the Immune synapse mask outline mask, and plotted using GraphPad Prism. For display purposes only, raw 3D stacks were reduced to a single plane using a maximum projection transformation and subjected to a linear scaling of their intensity, identically across all conditions, for optimal visualization.

#### Preparation of Stimulatory Surfaces for Microscopy and T cell activation assays

For microscopy, glass slides or 8 well chambers were coated with either ICAM-1 alone or anti-CD3 (clone Hit3a from Biolegend)/anti-CD28 (clone CD28.2 from Biolegend)/ICAM-1 (R&D systems) in PBS for 2hrs at 37°C to analyze T cell migration or synapse formation, respectively. Slides were washed 3X with PBS prior to use. To analyze T cell stimulation, proliferation, and cytokine secretion, 96 well MaxiSorp plates were coated with anti-CD28 with/without ICAM-1 (1 μg /mL each) and varying doses of anti-CD3 for 2 hours at 37°C, followed by three washes with PBS to remove unbound antibody.

#### T cell Imaging

To image the T cell F-actin network during immune synapse formation in real time, patient or control T cells were transfected with Lifeact -gfp and 48hrs later were washed and resuspended in Stimulation Buffer and allowed to spread on anti-CD3/28/ICAM-1 coated surfaces, and T cell synapses were imaged using a 1 micron-thick Z stack around the area of cell-glass contact using a Leica SP8 at the NIAID microscopy core facility. To image T cells migrating randomly on immobilized ICAM-1 cells were resuspended in L-15 media (Lonza) supplemented with 6mM Glucose and allowed to adhere to ICAM-1 coated chambers. Cells were then imaged with DIC and either a 63X objective with a 500msec frame rate to visualize the dynamics of the leading edge of migrating cells or a 5X objective at a frame rate of 30sec to measure cell migratory behavior over a longer period of time and distance. Images were collected on a Zeiss Axiovert 200 M inverted microscope equipped with an MS-2000 automatic stage (Applied Scientific Instruments) and a 37°C environmental chamber using Slidebook (Intelligent Imaging Innovation). For fixed cell imaging of the actin network patient or control T cell blasts were allowed to spread on ICAM-1 (1 μg/mL) or anti-CD3/28/ICAM1 (1 μg/mL each) coated (1 μg/mL each) surfaces. Cells were allowed to spread or adhere for 10minutes prior to washing away of loosely bound cells in prewarmed stimulation buffer and fixation in 3% PFA-PBS. Fixation was quenched with 50 mM NH_4_Cl and cells were then permeabilized with 0.2% Triton X-100 in PBS. Following extensive washing and blocking in 2% BSA/ 1 mM EDTA in PBS, cells were stained with Phalloidin-AF647 for 30 minutes at RT, washed and mounted with Fluoromount-G (Southern Biotech) prior to imaging as before.

#### *Ex vivo* T cell Activation Assays

To analyze T cell CD25/CD69 upregulation as well as T cell proliferation, Naïve CD4^+^ T cells (for Pt comparisons) or pan-CD4 T cells (for knockdown experiments) were first isolated from PBMCs and then labelled with CellTrace Violet (5 μM for 20 minutes). Cells were then resuspended to 1×10∧6 cells/mL in complete RPMI and added to prepared maxisorp plates, 20-50,000 cells/well, technical replicates were included if enough cells were present). For CD25 and CD69 expression, cells were collected at 36hrs post activation, stained on ice with anti-CD25(PE) and anti-CD69(FITC) (Biolegend) for 45-60 minutes, washed repeatedly and analyzed by flow cytometry. For proliferation, cells were collected on day 5 or day 6 post activation, washed and resuspended in FACS buffer. Immediately prior to collection cells were stained with propidium iodide (PI) to mark dead cells. Populations were gated first on Live cells by PI exclusion, then by forward and side scatter followed by exclusion of doublets by FSC-A by FSC-H. For cytokine secretion T cell blasts, activated 7-21 days prior, were rested for 3hrs in basal RPMI and resuspended in X-vivo15 without IL2. To analyze secretion by TCR stimulation cells were added to prepared maxisorp plates with the indicated immobilized stimuli (10-50×10^4^ cells/well). Supernatants were collected following an 18-24hr stimulation. For stimulation by CD3/28 beads, PMA/I, or IL-2 cells were added to an uncoated 96-well tissue culture plate and the appropriate soluble stimuli were added for 18-24hrs prior to supernatant collection. For CD3/28 bead stimulation, stimulatory beads (Invitrogen) were added to cells at a ratio of 1:2, and mixtures were briefly centrifuged to encourage bead/cell interactions. PMA/I were added to a final concentration of 10 ng and 1 μg/mL, respectively. For IL-2 stimulation, cells were plated and then 2X IL-2 containing X-vivo15 media was added for the indicated final concentration. Supernatants were analyzed by using either the Legendplex Multi-Analyte Flow Assay Kit for human CD8/NK cells or the human Th cytokine panel (Biolegend).

#### ICAM-1 Adhesion Assays

To analyze inside-out integrin activation T cell blasts were washed 3X in PBS (with Ca^2+^/Mg^2+^) and resuspended in stimulation buffer in the presence of 1μg/mL AlexaFluor 647 conjugated F(ab’)2 goat anti-human Fc antibody (Jackson ImmunoResearch) with or without 2 μg/mL ICAM-1. Cells were left alone or stimulated with PMA(10ng/mL)/Ionomycin(1 μg/mL) and fixed after 10minutes with a final concentration of 3% PFA. As a positive control for ICAM-1 binding T cells were stripped of Calcium and Magnesium with 10mM EDTA buffer and resuspended in Stim Buffer containing 1mM Mn^2+^ supplementation in the place of Ca^2+^ and Mg^2+^. Cells were incubated with ICAM-1 and the AlexaFluor 647 conjugated F(ab’)2 goat anti-human Fc antibody and fixed as before. ICAM-1 binding was measured by flow cytometry.

To analyze outside-in integrin activation, ICAM-1 was immobilized on maxisorp plates by first coating the plates overnight with 2 μg /mL anti-human Fc antibody (Jackson ImmunoResearch), washing 3X with PBS, and then incubating with the indicated dose of ICAM-1 at 37 C for 2hrs. The plate was washed with stimulation buffer and blocked in stimulation buffer for 2 hours. While the plate was being prepared, primary CD4^+^ or CD8^+^ T cells were collected and rested in basal RPMI for 2-3 hrs. Following the rest, cells were stained with 1 μM CFSE for 10 minutes at room temperature. CFSE was then quenched with FBS, and the cells were extensively washed in stimulation buffer to remove excess CFSE. Cells were placed on the ICAM-1 coated plate in triplicate, centrifuged for 2 minutes, and incubated at 37°C for 30minutes. The plate was washed gently with prewarmed Stim Buffer 2-4 times, and fluorescence was measured by on a plate reader. ICAM-1 adhesion was expressed as a percentage of the maximum adhesion of control cells.

#### Analysis of T cell signaling events

Relevant T cell blasts were removed from stimulatory beads at least 48hrs prior to reactivation and resuspended in fresh IL-2 containing media. Prior to the stimulation, dead cells were removed with the live/dead removal kit (Miltenyi) and cells were washed extensively with basal RPMI and rested at 37 C in basal media for at 3 hrs in basal media. Cells were then resuspended in either basal media or stimulation buffer. For analysis of early signaling events by western blot, resuspended cells were stimulated with anti-CD3/28 (1 μg/mL each) and crosslinked with Protein A (0.2 μg/mL) for the indicated times. After the indicated time, cells were lysed in TritonX-100 lysis buffer and kept on ice until the end of the time course. Lysates were then cleared by centrifugation (14,000RPM for 10 minutes), added to 4X reducing sample buffer and heated for 5mintues at 95°C. Samples were then analyzed by immunoblotting. For flow cytometry, cells were stimulated with the indicated crosslinked stimuli for the indicated time. Samples were then immediately fixed for 20miinutes with 3% PFA (final concentration), followed by quenching of the fixative, permeabilization for 30minutes with ice-cold methanol, and blocking with FACS buffer (PBS/ 2% FBS/ 1 mM EDTA). Cells were then stained for 1-18 hours for the indicated phosphorylated proteins, washed 3x with FACS buffer, and analyzed by flow cytometry. To analyze T cell calcium fluxes, T cell blasts from patienst or normal controls were collected, washed and rested in basal RPMI, and then loaded with 0.5 μM Indo-1 ratiometric calcium sensor (Thermo Fishher) in Powerload buffer (Thermo Fisher) for 20 minutes at room temperature followed by extensive washing to remove excess Indo-1. Cells were resuspended in Stimulation Buffer lacking extracellular calcium or FBS. Cells were then analyzed for calcium responses following TCR stimulation (10 seconds collection, addition of anti-CD3/28 antibodies, 20 seconds collection, addition of protein A crosslinker, 90 seconds collection, addition of 1 μM extracellular calcium, collect for total of five minutes) or following chemically induced release of ER Ca^2+^ stores (30 seconds of collection, addition of 1 μM thapsigargin, 90 seconds collection, addition of 1 μM extracellular calcium, collect for total of five minutes). Calcium level was expressed as a ratio of fluorescent spectrum of the Indo-1 dye in the bound (396 nm) to unbound state (737 nm).

#### Analysis of IL-2 induced Proliferation, CD25 and CD132 expression, and pSTAT3/5 activation

To analyze T cell proliferation downstream of the IL-2 receptor, Patient or Normal Control CD4^+^ T cell blasts were rested for 36 hrs in complete RPMI in the absence of IL-2. Cells were then washed, labelled with 1 μM CFSE for 5minutes at room temperature followed by quenching for 20 minutes with FBS and extensive washing, and dead cells were removed with the dead cell removal kit (Miltenyi. Cells were resuspended in complete RPMI with no IL-2, and plated at 2-5×10^4^ cells/ well in a 96 well plate. IL-2 containing media was then added to the indicated final concentrations and cells were incubated at 37°C for 5 days, after which CFSE dilution was measured by flow cytometry. To analyze CD25/CD132 expression, cells were prepared as above without CFSE labeling. 100 I.U./mL IL-2 was added to rested cells, and at the indicated times, cells were taken and fixed in PFA for 20 minutes. Following the completion of the time course cell pellets were resuspended in FACS buffer and stained with anti-CD25 (Alexa Fluor 488) and anti-CD132 (PE) for 30 minutes, washed 3X in FACS buffer, and analyzed by flow cytometry. To measure pSTAT3/5 phosphorylation, patient and control T cell blasts previously transduced by lentivirus were collected, washed 3X in basal RPMI, and rested for 3hrs in basal RPMI. Cells were then resuspended in basal RPMI at 1×10^6^/mL and cells were plated in a 96well V-bottom plate and warmed to 37°C. IL-2 at the indicated concentrations was then added for 10 minutes, and cells were fixed in PFA for 20 minutes and permeabilized in ice-cold methanol for 30 minutes followed by extensive washing in FACS buffer. Cells were then stained for pSTAT3 and pSTAT5 for 1 hr at RT and analyzed by flow cytometry.

#### Fas-and TCR mediated cell death

To determine sensitivity of patient cells to Fas-Mediated apoptosis, patient or control PanT cell blasts were collected and resuspended to 1×10^6^ cells/mL in fresh IL-2 containing complete RPMI. Between 0.5-1×10^5^ T cell blasts were added in triplicate to a 96 well flat bottom tissue culture plate. Crosslinked anti-Fas antibody (clone APO-1-1, Thermo Fischer) was prepared by mixing 10 μg antibody with 2 μg Protein A for 20 minutes at room temperature. Crosslinked anti-Fas was then added to cells at the indicated concentrations, and the cells were incubated at 37°C for 18 hrs. Cell death was measured by Propidium Iodide exclusion by flow cytometry and values were expressed as the percentage of dead cells (PI positive)/ total. To determine TCR restimulation-induced cell death, stimulatory plates were prepared as described above, and 0.5-1×10^5^ T cell blasts resuspended in IL-2 containing media were added in triplicate. Cells were allowed to reactivate for 18-24 hours followed by repeated pipetting and transfer to FACS tubes and analysis by flow cytometry. PI was added immediately prior to sample collection, and cell death was expressed as a percentage of dead cells/ total.

#### Neutrophil chemotaxis

Neutrophil chemotaxis was measured using EZ-TAXIScan instrumentation (Effector Cell Institute, Tokyo, Japan) which monitors the migration of PMN across a 260 µm platform separating the “Cell” well from the “Chemoattractant” well. Isolated neutrophils (5×10^3^ cells in 1.0 µL) were added to the “Cell” well of the EZ-TAXIScan and 1.0 µL of either buffer, fMLF (5×10^−8^ M), or C5a (5×10^-8^ M) was added to the opposing “Chemoattractant” well. Digital images of the migrating PMNs were captured every 30 sec for 1 hour.

Images were converted to stacks using the ImageJ software (version 1.46r; NIH). Ten randomly selected cells were electronically traced using the ImageJ plug-in, MTrackJ, and the sequential positional coordinates of individual migrating cells were determined as a function of time. The tracks of individual migrating cells were reconstructed and plotted with the position of each cell anchored at the origin at *t* = 0. Since data were collected with time and position, multiple parameters could be derived – overall distance, directed distance (parallel to the chemoattractant, random distance (orthogonal to the chemoattractant), overall velocity, directed and random velocity vectors, and time-to-event analysis (number of cells completing migration and elapsed time).

A modified Boyden chamber chemotaxis chamber was used to asses additional chemoattractants and doses. Isolated neutrophils (1×10^7^ cells/mL HBSS without divalent cations) were incubated with the cell-permeant fluorescent probe, calcein-AM (5 µg / 5 µl of DMSO/ml of cell suspension) for 15 min in a 37°C incubator wrapped in foil to protect from light. At the end of the incubation, the cells were washed, counted, and resuspended at 3×10^6^/ml in 2% BSA/HBSS with divalent cations. The chemoattractants were diluted in 0.1% HSA in PBS and 29 µl of each chemoattractant was added to the bottom well of a 96-well ChemoTx® Disposable Chemotaxis System chamber that is fitted with a framed polycarbonate filter 6-10

µm in depth with 5.0 µm pores at a density of 4×10^5^ pores/cm^2^ (Neuro Probe, Inc., Gaithersburg, MD). The filter frame was replaced on the 96-well plate and 25 ul (75,000 cells) were pipetted onto an 8 mm^2^ filter spot surrounded by a hydrophobic mask, forming a hemispherical droplet directly above the chemoattractant well. The chamber was incubated for 45 min at 37°C, then the remaining cells were aspirated from atop the filter. EDTA (15 µl of 2 mM solution) was added to each filter spot and incubated additional 30 min at 4°C to promote cell detachment from the filter. The EDTA was aspirated from the filter and the chamber centrifuged for 5 min at 1000 rpm. The fluorescence of each well was determined using a fluorescent plate reader (Gemini EM, Molecular Devices, San Jose, Ca) using a λ_excitation_ = 490 nm and λ_emission_ = 520 nm. Fluorescence was converted to number of migrating cells using a standard curve of calcein-AM loaded cells.

#### Immunoblotting

To prepare lysates, cell pellets were either immediately lysed in 1% Triton X-100 buffer, 0.5% NP-40 buffer, or flash freeze-thawed five times before 25-minute incubation in CHAPS lysis buffer (used for co-IP samples to maintain low affinity interactions). Jurkat and primary T cells were lysed in 1% Triton X-100 (150 mM sodium chloride, 1% Triton X-100, 50mM Tris pH 8.0 with protease and phosphatase inhibitor tablets (Roche)) when analyzing T cell signaling or 0.3% CHAPS buffer (0.3% CHAPS w/v, 40 mM HEPES pH 7.5, 120 mM NaCl, 1mM EDTA with protease inhibitor tablet (Roche)) for Co-IP experiments. To analyze WRC interaction in overexpression experiments, HEK 293T cells were lysed in 0.5% NP40 lysis buffer (0.5% NP40 w/v, 150mM NaCl, 50mM Tris-Cl with protease inhibitor tablet (Roche)). Following cell lysis for 15-25 minutes on ice, lysates were cleared by 10-15 minutes of centrifugation at 14,000 RPM at 4°C. Protein concentration of clarified lysates was determined using the Micro BCA Protein Assay Kit (Thermo Fisher). Samples were denatured in reducing (10% BME) NuPAGE LDS sample buffer (Thermo Fisher), boiled for 5 minutes at 95°C, and separated on 4-12% Bis-Tris gels (Invitrogen). Proteins were transferred to nitrocellulose membranes and blocked with 5% milk. Incubation with primary antibodies occurred for 2 hours at room temperature or overnight at 4°C, followed by 1-hour incubation with goat anti-mouse AlexaFluor 680 or goat anti-rabbit AlexaFluor 800 secondary antibodies (Thermo Fisher). Alternatively, HRP-conjugated anti-mouse or anti-rabbit secondary (Southern Biotech) were used to visualize proteins by chemiluminescence. Membranes were imaged on the Azure c500 Imaging System (Azure Biosystems) or exposed to autoradiography film and ImageJ software was used for all image analyses.

#### Immunoprecipitation

Lysates were prepared as described above. Clarified lysates (800-1000 μg) were incubated with primary antibody at 4°C for 1 hour, then for an additional 2 hours with magnetic Dynabeads Protein A (Thermo Fisher). Protein complexes were isolated, washed 3-5 times in lysis buffer, and immunoblotted as described or used in Mass-Spec analysis as described below.

#### Recombinant protein purification and biochemical assays

Recombinant human WRC containing full-length (FL) Cyfip1, FL Hem2, FL HSPC300, Abi2 (1-158) and WAVE1 (1-230)-(GGS)_6_-(485-559), which we named WRC230VCA, was purified as previously described.(*36*) The WRC containing M373V Hem2 behaved identically to the WT during expression and various chromatographic steps in purification, suggesting M373V did not affect folding, stability or WRC assembly. Other proteins, including the Arp2/3 complex, actin, WAVE1 VCA, GST-Tev-Rac1 (Q61L/P29S, 1-188), and Rac1 (Q61L/P29S, 1-188) were purified as previously described.(*37*) GST-thrombin-Arf1 (18-181) construct was obtained from Neal Alto’s lab and was expressed and purified by following a similar protocol used for GST-Tev-Rac1.(*37*) Untagged Arf1 (18-181) was obtained from MBP-3C-Arf1 (18-181) after HRV 3C protease cleavage to remove the MBP tag, followed by SOURCE 15Q anion exchange and Superdex75 size exclusion chromatography (GE Healthcare). To obtain a heterodimer of Rac1 (Q61L/P29S, 1-188) and Arf1 (18-181) that we use in the actin polymerization assay, we inserted a heterodimeric GCN4-derived coiled coil (CC) followed by a flexible GGS linker between the MBP-Tev-tag and Rac1 (using the acidic CC) or Arf1 (using the basic CC).(*38*) The MBP-CC-GTPases were expressed and purified individually from BL21(DE3)^T1R^ cells (Sigma), combined in ∼1:1 molar ratio to form a heterodimer, treated with Tev to remove the MBP tag, and purified by SOURCE 15S cation exchange and Superdex75 size exclusion chromatography (GE Healthcare).

GST pull-down assays and pyrene-actin polymerization assays were performed by following previously established protocols.(*37*) Prior to use, Arf1 (alone or in heterodimer with Rac1) was loaded with GDP, GTP (in pull-down) or the unhydrolyzable GTP analog GMPPNP (in actin polymerization assay) by following a similar protocol used for Rac1 WT, except that we changed the incubation condition to 37°C for 45-60 min in order to maximize the loading efficiency for Arf1.(*37*) Note that Rac1 (Q61L/P29S, 1-188) is constitutively active and cannot be loaded with GDP under the same condition, despite various temperature and incubation time that we tested.

#### IP-Mass Spectrometry

To analyze Hem1 and Wave2 interacting proteins three sets of IP-s were done followed by quantitative mass spectrometry. Firstly, from 1×10^8^ cells, an immunoprecipitation was performed with either a Rabbit Isotype (CST) or Wave2 antibody (CST) followed by two washes in lysis buffer and two more in detergent-free lysis buffer. Magnetic IP beads were then resuspended in sodium dodecanoate and boiled. The second set of IPs was performed using the Wave2 antibody in either lysates generated from 5×10^7^ Wave2-deficient Jurkat cells, Hem1-WT-Flag reconstituted cells, or Hem1-M371V-Flag reconstituted cells. Samples were again washed 2X in lysis buffer and 2X in detergent free lysis buffer and resuspended in sodium dodecanoate and boiled. For the third set of IPs, FLAG was immunoprecipitated from 5×10^7^ Hem1-deficient cells, Hem1-WT-Flag reconstituted cells, or Hem1-M371V reconstituted cells. Samples were again washed 2X in lysis buffer and 2X in detergent free lysis buffer and resuspended in sodium dodecanoate and boiled for 5 minutes.

Briefly, each control/experiment pairs were compared essentially by LC/MS/MS using the affinity enrichment approach with isotope-based quantitation based on post digestion reductive di-methylation and data processing using MaxQuant. (*39–41*) More specifically, to each sample (containing 100 μL 1% sodium dodecanoate), 10 μL of a urea-unfolded solution of non-reduced Hen ovalbumin (about 0.3 picomole) was added. To ensure a controlled pH, 50 μL of 1M Tris (pH 8.0) was added and then 3 μL of 0.4M DTT was added and the sample heated to 50°C for 20 minutes. After cooling to room temperature, 6 μL of 0.5 M chloroacetamide was added and the samples incubated in the dark for 1 hour, after which 3 μL of beta-mercaptoethanol was added to scavenge remaining chloroacetamide. Next the samples were diluted to 0.1% dodecanoate and 2 μg of trypsin (5uL, V5113), mixed thoroughly, and incubated overnight at room temperature. Sodium dodecanoate/laurate was removed by adding 50 μL of 10% formic acid, samples were washed three times with 0.5 volumes ethyl acetate, and phase separation was reestablished after vigorous mixing and a short spin at 800xg.(*42*) After extraction, the aqueous fraction was moved to a fresh vial and a further 200 μL of 0.4% formic acid was added to recover residual sample from the leftover aqueous phase in the original sample vial. This solution was withdrawn and combined with the bulk of each sample. Each sample was dried down briefly with a stream of warm nitrogen gas before being chilled in ice water. Each sample was then applied to STAGE tips and labeled by reductive dimethylation on column essentially as described as protocol C.(*40, 43*) After elution, most of each sample was mixed with its pair partner. Light labeling (+28) was used for the control and intermediate (+32) was used for the pull-down samples. To improve the quality of ratio estimation with high background samples from lightly washed pull-downs, the mixed sample was fractionated using a concatenated high pH (pH 10) reversed phase approach as described. (*41*) Future applications of high pH fractionation of dimethyl labeled samples such as this might be improved by performing them at slightly lower pH to avoid the potential beta-effects of isotopic forms at higher pH 10. (*42*) This effect may have played a role in a small number of declinations by software to make ratio calls in later data analysis.

Data was next collected on an LC/MS/MS system comprising a Thermo nLC-1000 and Thermo Orbitrap Fusion Lumos configured with an EASY-Spray source. The nLC-100 used an A-solvent of 0.1% formic acid and a B-solvent of 93.75% acetonitrile 0.1% formic acid. From a final 40 μL sample, 18 μL was injected and loaded at 450 bar onto an EASY-Spray column (50 cm x 75 μm ID, PepMap RSLC C18, 2 μm) and separated at a flow rate of 100 nl/minute with a gradient of 5 to 22% in 215 minutes, followed by a gradient to 32% in 30 minutes, and then to 95% in 20 minutes. Parent spectra were collected using the orbit rap with a 120K resolution setting after which ion trap spectra of HCD fragmented targets were collected for up to 5 seconds. Peptide like mono-isotopic peak detection, 1E3 intensity, 2-7 charge state, and 2 over 2 second dynamic exclusion (20 second duration) were used as filters. Raw data were processed using MaxQuant 1.6.3.3 using a two-channel analysis with re-quantification and otherwise default settings.

#### Flow cytometry

Cell-surface receptor staining were performed on ice prior to fixation for at room temperature for 30 minutes – 18hrs following fixation/permeabilization. For intracellular staining, cells were fixed and permeabilized 3% PFA and 100% ice-cold methanol, washed at least 3X, and incubated with antibodies for 1-18 hours at room temperature. Flow cytometric data were acquired on a BD Fortessa with 18-color configuration, an LSRII, or a FACS Caliber and analyzed using FlowJo version 9.9.6 or 10.4.2.

#### Statistical Analysis

All statistical analyses were done in GraphPad Prism 8. * P<.05, ** P<.01, *** P<.0001. Differences that did not reach statistical significance are not indicated. For statistical tests done between normal donors and patient cells (when three or more patients were tested for that assay), a student’s t-test assuming unequal variance was performed. For knockdown and inhibitor experiments, where samples were compared between treated and untreated conditions, a nonparametric paired t-test (Wilcoxon matched-pairs signed-rank test) was performed.

### Materials

#### Antibodies

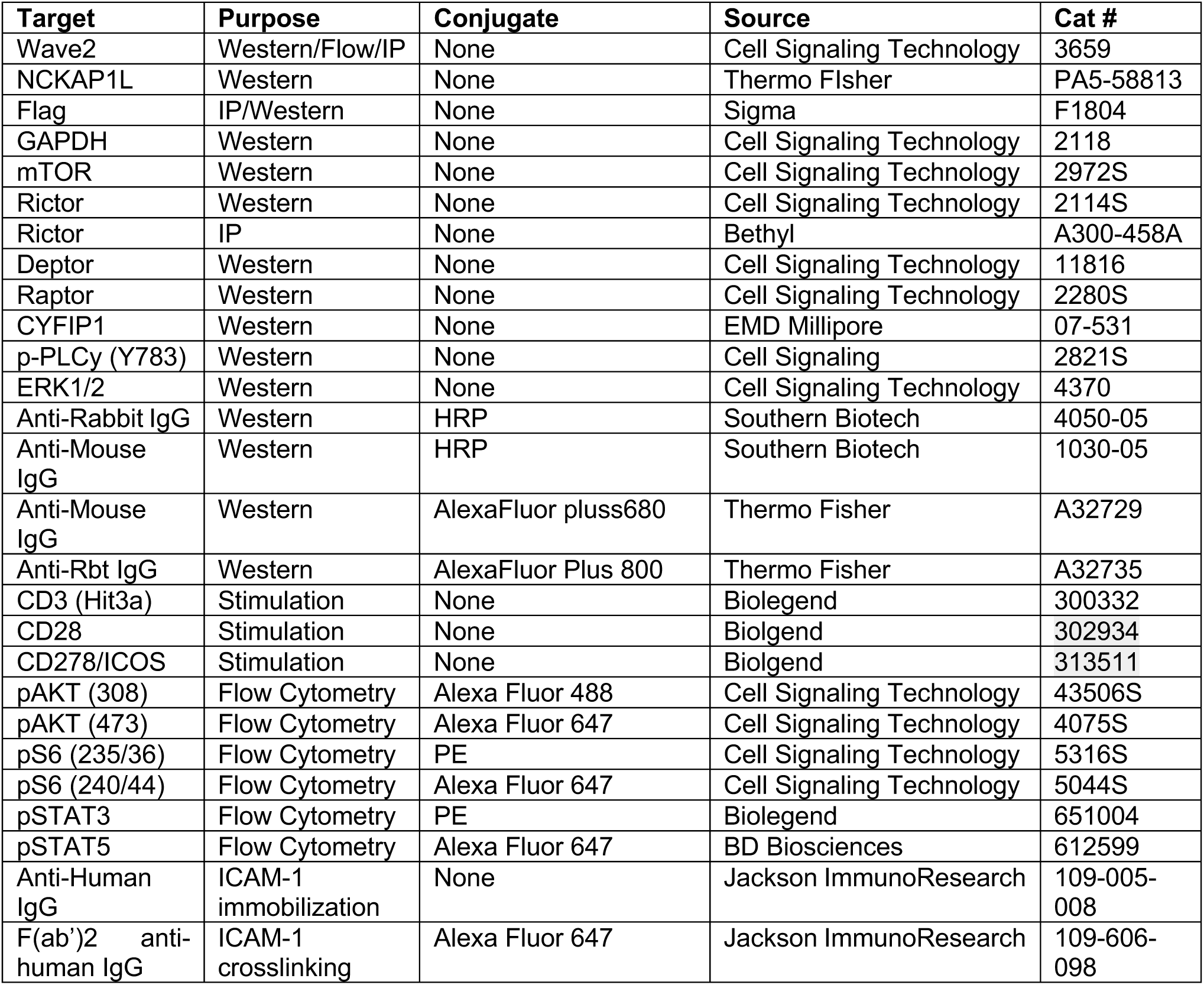

## Supplementary Text

### Patient Narratives

#### Pt 1.1

Pt 1.1 is an 11-year-old female born to non-consanguineous parents both from a Canadian First Nations/Ojibway ancestry. Whole exome sequencing (WES) revealed that the patient has three regions of absence of heterozygosity with an identity by descent of 0.86%, indicating the parents may be distant cousins. Following an uncomplicated pregnancy and healthy birth the patient quickly developed chronic rhinorrhea and cough during the first months of life. At 9 months of age the patient developed an upper respiratory tract infection with fever and a bulging anterior fontanelle and was diagnosed with viral meningitis. At the age of 15 months the patient was taken to the emergency room with increased work-of-breathing, cough, fever, lethargy, decreased oral intake, and diarrhea. A chest X-ray showed right middle and lower lobe infiltrates with *Streptococcus pneumoniae* identified in blood cultures. She developed necrotizing pneumonia with a pleural effusion, requiring intubation and chest tube placement and hydrocephalus with presumptive treatment for meningitis. During this visit the patient was noted to have pulmonary hypertension, hepatomegaly, and splenomegaly which resolved during treatment. The patient was treated with vancomycin and 3^rd^-generation cephalosporin and a 2-week course of intravenous ampicillin/ceftriaxone. Subsequently, the patient suffered from two additional rounds of pneumoniae with cough, fever, and respiratory distress within the first two years of life. She was diagnosed with asthma with nocturnal hypoxia at 15 months-of-age, which required night-time O_2_ supplementation with limited benefit from inhaler treatment (including fluticasone-salmeterol and albuterol). Chronic middle ear infection led to surgery for tympanostomy tubes at 4 years-of-age. The patient subsequently developed increased exercise-intolerance, cough/tachypnea, and work of breathing, culminating in hospital admission at 6 years-of-age, where pulmonary function tests showed a restrictive lung pattern (FEV1 = 36% predicted, FVC = 39% predicted, FEV1/FVC =0.83) and a CT scan demonstrated diffuse interstitial infiltrates. Further analysis revealed splenomegaly (8 cm below left costal margin), mild hepatomegaly (3 cm under right costal margin), and diffuse generalized lymphadenopathy. Immunophenotyping revealed elevated IgG and IgE levels (23.1 g/L and 2000 IU/mL, respectively) with normal IgM and IgA levels, normal or elevated double-negative CD3^+^ T cells (3.4% at one timepoint and 8%-11% at another), and normal vitamin B12 concentrations. Biopsy of the axillary lymph node demonstrated an encapsulated lymph node with numerous reactive germinal centers and a reactive paracortex and no double-negative T cells found; it was diagnosed as reactive follicular hyperplasia. Lung biopsy of the right middle lobe demonstrated chronic interstitial lung disease, and moderate to severe diffuse interstitial pneumonitis of predominantly lymphocytic origin with increased alveolar macrophages and rare eosinophils and neutrophils. The biopsy also revealed mild bronchiolitis, mild pulmonary arterial medial hypertrophy, and pleural fibrosis and hemorrhage. A next generation gene mutation panel for autoimmune lymphoproliferative syndrome (ALPS) and ALPS-related genes, including the CASP8, CASP10, FADD, FAS, FASLG, ITK, KRAS, NRAS, and MAGT1, showed no mutations in these targets. The patient was started on steroids and sirolimus, which improved or resolved the patient’s respiratory difficulty and lymphadenopathy/splenomegaly. At 7 years-of-age the patient was readmitted for one week with fever and chills, left-sided knee pain, and positive blood cultures for *Streptococcus pneumonia*. She was treated with ceftriaxone and amoxicillin. Ultrasound of the knee showed a small fluid collection with a thickened bursa that persisted following discharge. Since then, the patient has received amoxicillin prophylaxis that has helped control subsequent infections.

Following initial improvement on sirolimus and steroids the patient’s respiratory state diminished after she was weaned off of steroid treatment, where-upon the patient developed worsening cough and exertional dyspnea. At chest CT scan demonstrated diffuse ground-glass appearance in both lungs with septal thickening and areas of air-trapping, subsegmental atelectasis and bronchiectasis. The dose of sirolimus was increased, and steroids were restarted with minimal benefit. The patient was later switched to mycophenolate mofetil (MMF) with secession of sirolimus/steroids to control the lymphoproliferative disease. Currently the patient remains largely symptom free on MMF.

Additionally, the patient has a history of recurrent methicillin-resistant *Staphylococcus aureus* skin pustules and eyelid hordeolum/blepharitis, as well as mild atopic dermatitis. She has been treated with once-weekly bleach baths. The patient has no history of herpes simplex virus (HSV), molluscum, warts, or urticaria.

In addition to the patient’s immunological deficiency, she also exhibits signs of a mild extra-hematopoietic disease. Anatomically, the patient has pectus carinatum and has poor dentition with small, worn down central and lateral incisors. Echocardiogram demonstrated trivial tricuspid valve deficiency. The contribution of other rare homozygous variants to the patient clinical phenotype cannot be excluded as she has several homozygous variants of unknown clinical significance that segregate with disease (Table S3).

#### Pt. 2.1

Pt 2.1 is a 16-year-old female born to non-consanguineous parents of Mexican origin. Family history was negative for lymphoproliferation or immune deficiency, but she had a brother that died at 2.5 years of age due to infections and renal disease, no tissue was available for genetic screening of this sibling. The deceased brother had a positive PPD test and received isoniazid for 9 months; no more history is available for this sibling. Pt 2.1 was born healthy at term. At six months of age, she started having more than 6 episodes of upper respiratory infections and recurrent ear infections a year and pulmonary infections accompanied by wheezing for which she received asthma treatment with inhaled steroids and albuterol. Antibiotics were given during exacerbations. After age 2, she continued to have recurrent pneumonias and was hospitalized once due to this diagnosis. Also, after 2 years of age, she developed failure to thrive with height and weight below the third percentile for each; multiple sweat tests were performed and were normal. She was diagnosed with celiac disease and selective IgA deficiency during this period of time. She was first evaluated by Immunology at eight years of age and found to have clubbing of the fingers, and a purulent ear infection. Her immunologic evaluation showed a high IgG level and undetectable IgA and IgM. T, B, and NK cell numbers were within normal ranges, but NK cell function was decreased. The patient has had recurrent lymphadenopathy and splenomegaly; however, the percentage of *αβ* double negative T cells was below 2%, and the rest of the ALPS evaluation was negative. A chest CT scan revealed peribronchial thickening and bronchiectasis as well as hilar adenopathy. In consideration of her chronic pulmonary symptoms and bronchiectasis, she was started on immunoglobulin replacement and prophylactic treatment with azithromycin. The patient has no history of HSV, molluscum, warts, or urticaria, and acute infection by cytomegalovirus (CMV) or Epstein-Barr virus (EBV) infection. She had one episode of cellulitis post pneumococcal vaccination. The patient has no dysmorphic features and no learning disabilities have been detected. Recently, at 15 years of age Patient 2.1 had an episode of acute abdominal pain during which she was diagnosed with acute cholecystitis and cholelithiasis requiring uncomplicated cholecystectomy.

#### Pt. 2.2

Pt 2.2 is the younger sibling of Pt 2.1 and is currently 10 years old. He was first evaluated at two years of age because of a history of recurrent ear infections that started at 12 months of age and recurrent fevers and two prior hospitalizations due to fever with undetermined diagnoses. At the time, he had elevated levels of IgG (1879 mg/dL), IgA (179 mg/dL) and IgM (151 mg/dL). Lymphocyte phenotyping showed increased B cells. At three years of age, he had fever and diffuse lymphadenopathy of the cervical, mediastinal, and mesenteric lymph nodes accompanied by hepato-splenomegaly (Supplemental images C). A biopsy of a cervical lymph node performed during this hospitalization showed reactive follicles with subcapsular sinus histiocytosis and emperipolesis suggestive of Rosai-Dorfman disease. Genetic testing for ALPS was negative, and double-negative T cells were not increased. During the same hospitalization, he was noted to have nephrotic range proteinuria and hematuria. In the context of this nephrotic syndrome, a biopsy was performed and showed glomerulonephritis with deposition of immune complexes in a subendothelial and mesangial pattern. The biopsy findings were compatible with lupus nephritis class IV-V but the patient did not fulfill other systemic lupus erythematosus (SLE) criteria at the time. In particular, ANA and anti-DNA auto antibodies were negative. He was started on steroids and MMF. He continued to have multiple episodes of respiratory infections, including condensing pneumonias and severe, purulent ear infections and his renal disease seemed to be exacerbated by these infections. A chest CT scan obtained in between pulmonary infections revealed bilateral ground glass opacities with no bronchiectasis (Supplemental images). At 7 years of age, he had an exacerbation of the nephrotic syndrome requiring pulsed steroids and initiation of rituximab. After treatment with rituximab, he was also started on subcutaneous immunoglobulin infusions to control recurrent infections. His renal disease has been partially controlled with MMF and rituximab but appears exacerbated by ongoing infections. Despite immunoglobulin replacement and antibiotic prophylaxis, he continues to have recurrent episodes of infection, including multiple episodes of pneumonia. One event was associated with pneumococcal bacteremia. He has recently had a pleural effusion. He also continues to have recurrent and persistent otitis media. Between pulmonary infections, he has chronic cough and wheezing. He was diagnosed with asthma at the age of 7 and started on inhaled steroids and bronchodilators, which has led to only partial control of symptoms. After the first episode of lymphadenopathy, he has had multiple exacerbations of lymph node enlargement and hepatosplenomegaly, approximately once a year. During these episodes, he shows enlargement of cervical, mediastinal, mesenteric and inguinal lymph nodes (Supplemental Figure S1A). Multiple biopsies performed during these episodes show reactive follicular enlargement and have ruled out malignancy. Rosai-Dorfman disease has been identified twice in cervical lymph node biopsies. The most recent chest CT scan shows persistence of supraclavicular adenopathy, mediastinal lymph node enlargement, as well as ground-glass and consolidative opacities within the right upper and lower lung lobes. The patient has no history of HSV, molluscum, warts or urticaria, and acute infection by CMV or EBV has not been observed. He has no dysmorphic features; he has problems in school apparently due to multiple absences due to his recurrent and extended periods of medical absence.

#### Pt 3.1

Patient 3.1 is 10-year-old male born to healthy first-degree cousins in the United Arab Emirates. Whole exome sequencing revealed that the patient has eight regions of absence of heterozygosity with an identity by descent of 4.6%, confirming the parental relation. Though born to healthy parents, the patient had three deceased elder brothers that died at 3 months of age from fever and organ failure indicative of sepsis along with dysregulated inflammation similar to HLH. At birth, the patient exhibited jaundice and required an extended hospital stay. At 13 months of age, he was hospitalized for invasive and disseminated HSV1 infection, manifesting with muco-cutaneous lesions, hepatitis, and encephalitis (with epilepsy). He required a 4-week hospitalization and anti-seizure therapy due to herpetic meningo-encephalitis. Long-term consequences of the infection included developmental delay and sensorineural hearing loss and lack of speech. At 18 months of age, the patient developed sudden hearing loss. His subsequent clinical history includes several instances of high fever and splenomegaly requiring hospitalization and administration of antibiotics with blood cultures testing positive for *Pseudomonas aeruginosa*, as-well-as persistent asthma, allergic rhinitis, and allergies (anaphylaxis) to food (shellfish). Additional infections include molluscum contagiosum lesions in the upper chest and head at 7 years, mucoid otitis media at 9 years, and roseola infantum at 10 years of age. Despite normal serum immunoglobulin levels at the time of testing, the patient had poor pneumococcal-specific antibody titer following polysaccharide pneumococcal vaccine administration and developed autoimmune thrombocytopenia following MMR vaccination. In addition to the immunological abnormalities, the patient is obese and, along with his elder sister, displays bleeding tendencies with gum and skin bruising. He has an abnormal platelet function test, low factor 12 level, and prolonged partial thromboplastin time. The family history is significant for easy fractures and recurrent fever in the oldest brother and temporal arachnoid cysts with history of craniotomy, mental developmental delay (autism-like), and a ganglion cyst excision (right wrist) in another brother. All siblings have significant dentition problems with retention of primary teeth. These symptoms may indicate the presence of one or more additional genetic diseases within the family. Supporting this likelihood, the patient has several homozygous variants of unknown significance, including in proteins known to be important for bone development and cognitive function (Table S3).

#### Pt 4.1

Patient 4.1 is a 2.5-year-old female born to non-consanguineous parents from north-Africa with two healthy sisters. At approximately one year of age, the patient developed pulmonary symptoms with alveolar interstitial syndrome and pancytopenia; however, no bacterial or viral infection was detected. One month later, she was admitted to the hospital for persistent fever, splenomegaly, hyperferritinemia, hypofibrinogenemia, hypertriglyceridemia, and an EBV viral load of 3.8 log copies/mL. The patient was subsequently diagnosed with EBV-driven hemophagocytic lymphohistiocytosis. No defect in T cell degranulation, perforin expression, or hair color was detected. She was treated with corticosteroids, cyclosporine, and two rounds of rituximab. Following an initial recovery, the EBV viral load again increased after 8 months to 3.6 log copies/mL with no detectable anti-EBNA IgG, indicating a lack of seroconversion and poor specific antibody production. She developed hepatosplenomegaly at one year of age, which continues to persist. The patient has also had recurrent otitis media and pneumonia with the development of bronchiectasis and recurrent asthma. Additionally, she has features of a syndromic disease with several present at birth, including intracerebral ventricular dilatation (unilateral) and ventricular heterotopia, liver calcification (liver hamartoma), kidney dilatation, cardiomegaly, and muscular ventricular septal defect with dilatation of the ascending aorta. Following birth, the patient developed aortic bicuspidia, and had oral feeding issues requiring a gastric tube.

**Fig. S1.**
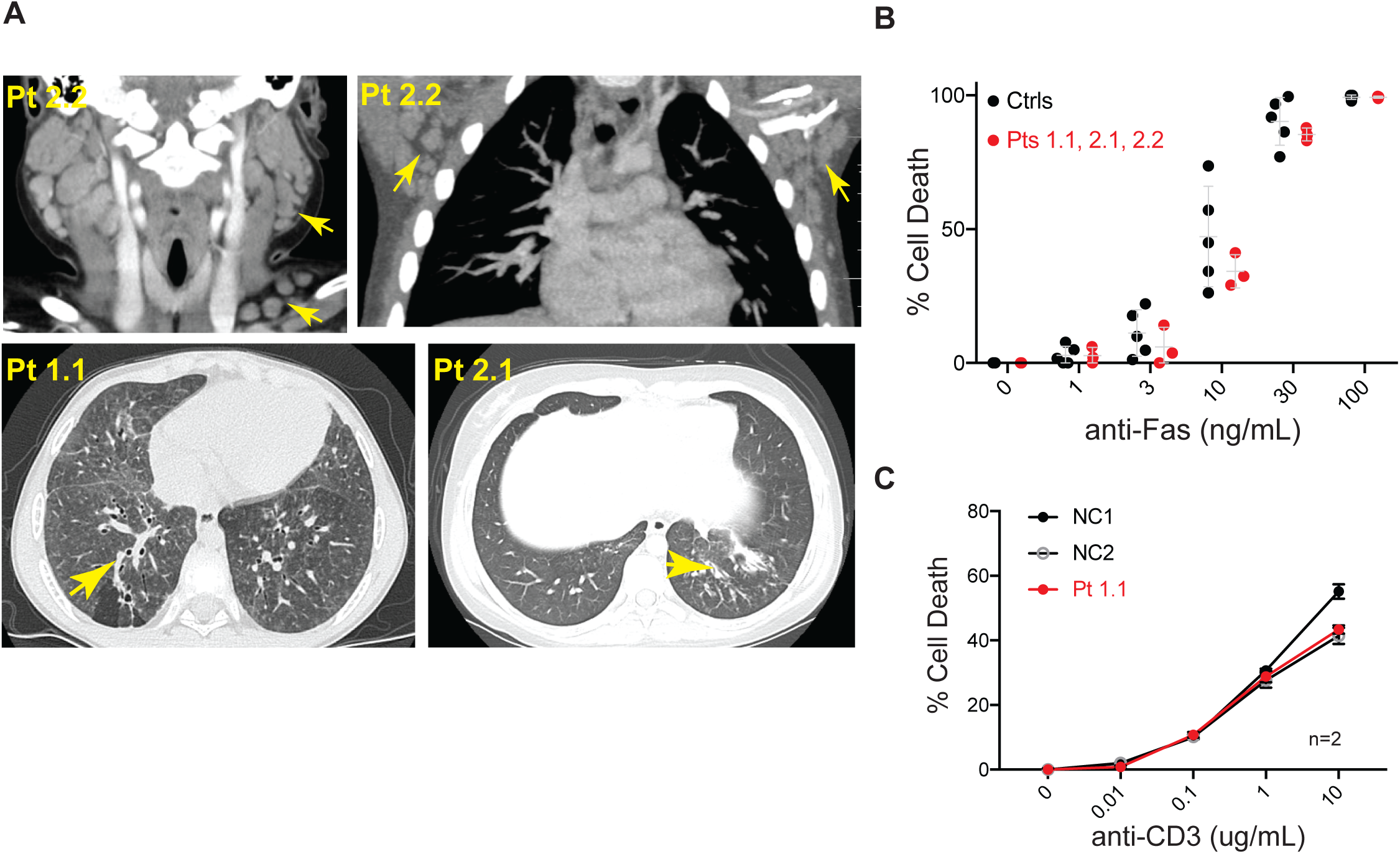
Patients develop bronchiectasis and lymphadenopathy that is not associated with a defect in Fas- and TCR-mediated apoptosis. (**A**) CT showing enlarged lymph nodes in Pt 2.2 (top) and bronchiectasis in Pts 1.1 and 2.1 (bottom) indicated by yellow arrows. (**B**) anti-Fas-mediated apoptosis determined by Propidium Iodide (PI) exclusion in CD8^+^ T cell blasts derived from patients and normal controls. (**C**) Cell death assessed by PI exclusion following a 24-hour stimulation on immobilized anti-CD3 at the indicated dose and anti-CD28/ICAM-1 (1 ug/mL) each.

**Fig. S2.**
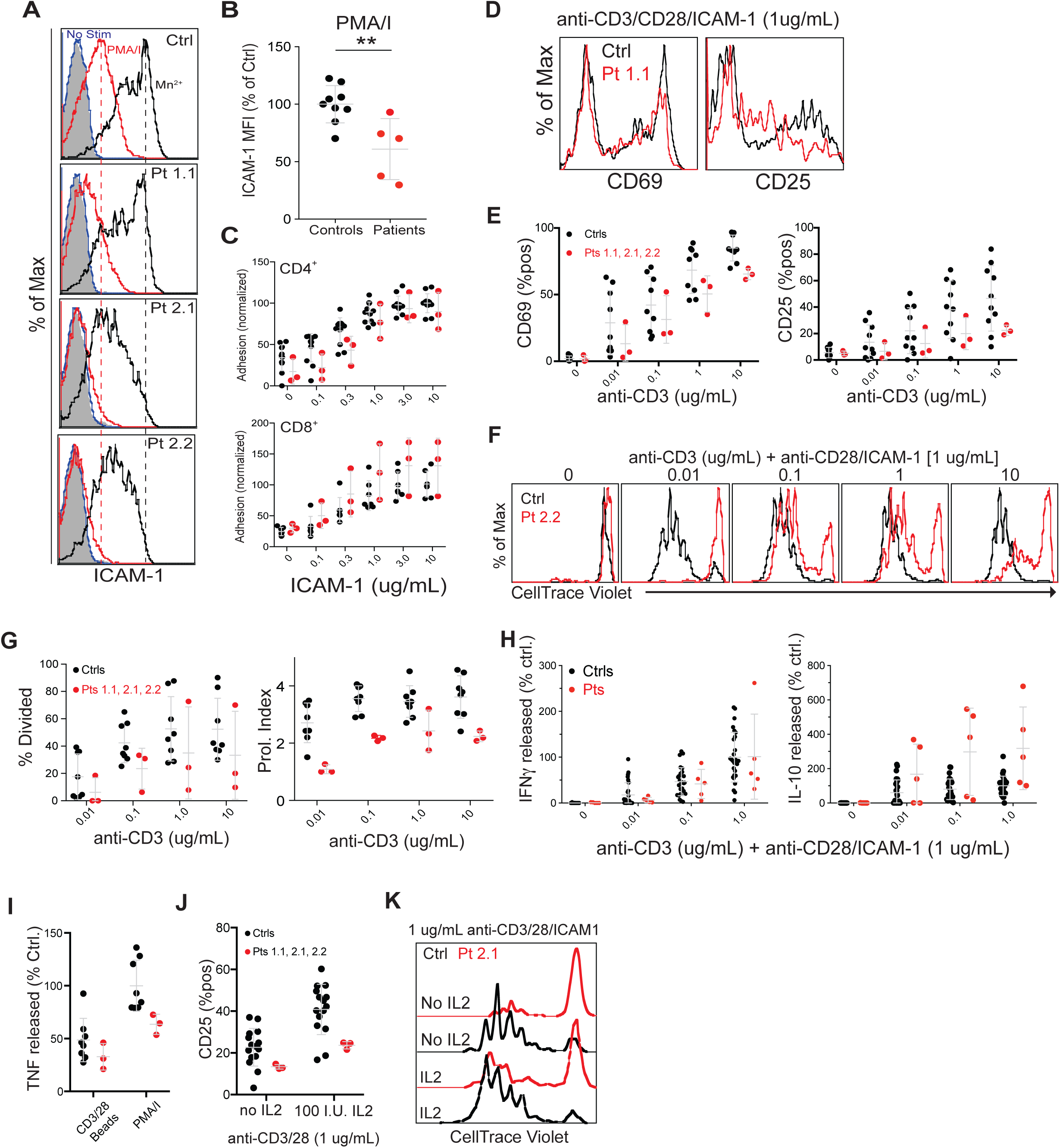
Hem1 deficient patient T cells are defective in primary activation and selective effector function. (**A**) Representative histograms of soluble ICAM-1 binding following no stimulation, 10 minutes of PMA and Ionomycin (PMA/I) stimulation, or Manganese-induced (Mn^2+^) integrin activation. (**B**) Averages of soluble ICAM-1 binding to PMA/I-stimulated cells normalized to the control average, as shown in (A). (**C**) CD4^+^ and CD8^+^ T cells adhering to the indicated dose of immobilized ICAM-1. Values were normalized to the average of the maximum control response. (**D**) Representative flow histograms of CD25 and CD69 expression on patient T cells following 36hr stimulation with anti-CD3, anti-CD28, and ICAM-1 (1 ug/mL each) coated surfaces. (**E**) Combined results of three independent experiments showing CD69 and CD25 upregulation over a range of CD3 stimulation with 1 ug/mL anti-CD28 and ICAM-1. (**F**) Representative flow histograms of patient T cell proliferation 5 days post-stimulation with the indicated doses of anti-CD3/28 and ICAM-1. (**G**) Quantification of the % divided (% of initial population that divided at least once) and the proliferation index (# of division/dividing cell) of patient cells and controls from three independent experiments. (**H**) IFN*γ* and IL-10 secreted from CD4^+^ T cell blasts following a 36hr re-stimulation with immobilized anti-CD3/28 and ICAM-1 at the indicated concentrations. (**I**) IL-2 and TNF secretion following a 36-hour restimulation with either anti-CD3/28 stimulatory beads or PMA/I. Numbers are normalized to the control response to PMA/I. (**J**) CD25 and (**K**) T cell proliferation measured by CellTrace violet dilution in the presence or absence of exogenously added IL-2 (100 I.U./mL).\*\**P ≤* 0.01.

**Figure S3.**
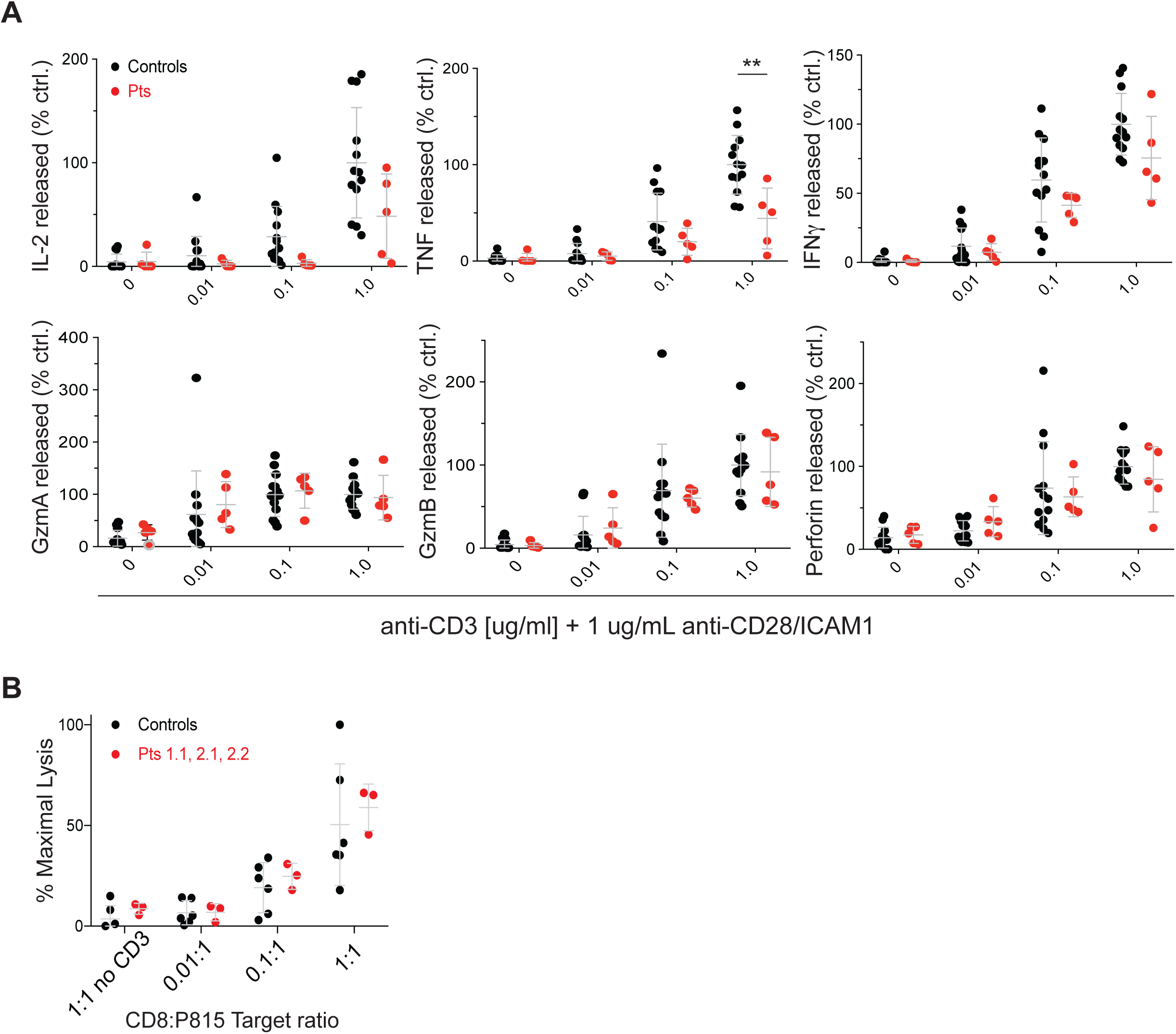
Patient CD8^+^ T cell cytokine production and cytolytic activity. (A) Cytokine release from Patient CD8^+^ T cells stimulated on the indicated dose of plate bound anti-CD3 and anti-CD28/ICAM1 [1 ug/mL] each. **(B)** Lysis of P815 target cells coated with 1 ug/mL anti-CD3 prior to incubation with CD8^+^ T cell blasts derived from either Patient or Normal Controls

**Figure S4.**
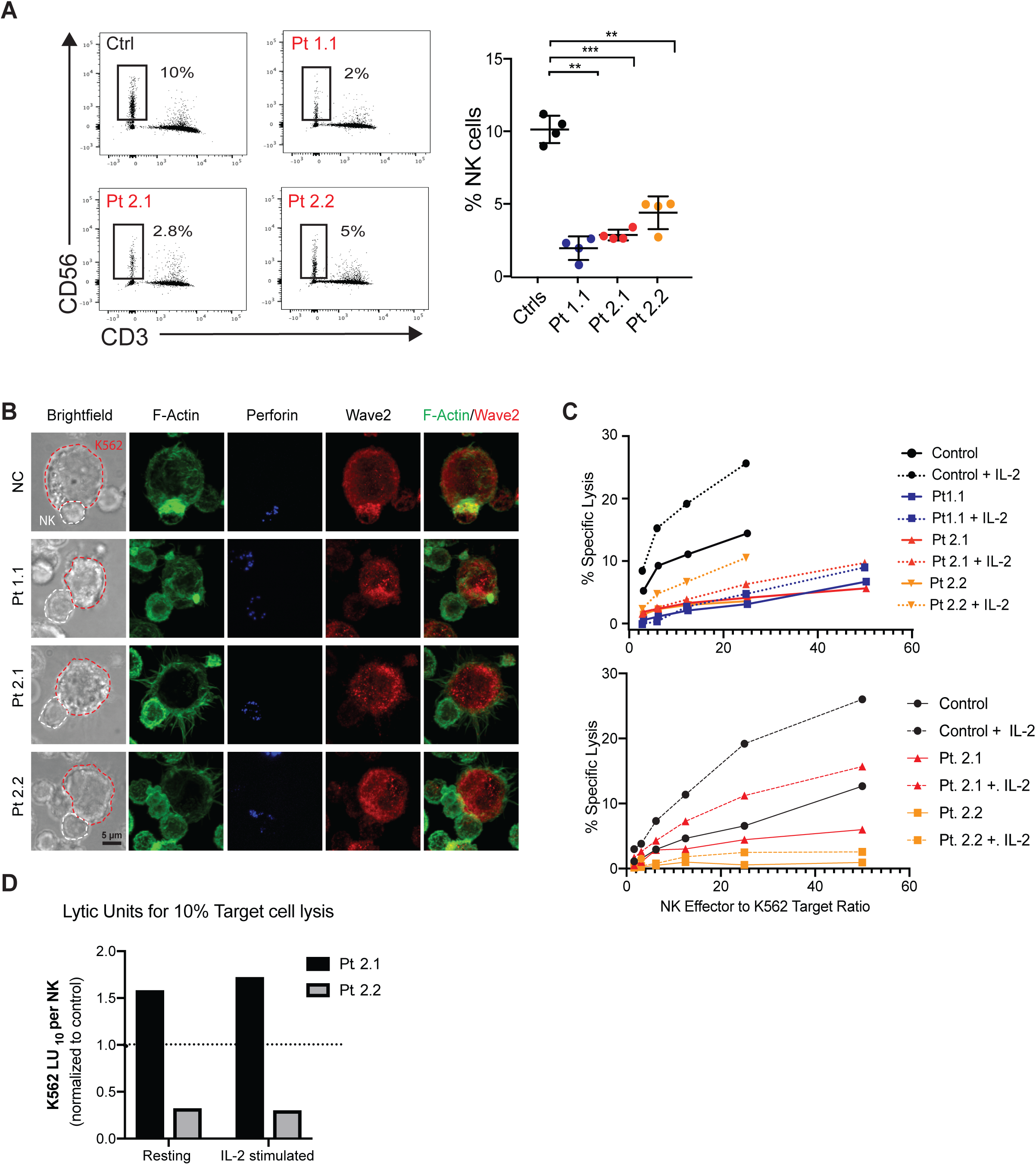
Hem1 patients have reduced NK cell numbers and diminished cytotoxicity associated with decreased actin accumulation at the NK cell immunological synapse. (**A**) Gating strategy (left) and percentage of NK cells in PBMCs from Hem1 patients and normal controls. (**B**) Representative imaging (expansion of Fig. 2I) of NK cell immunological synapses for patients 1.1, 2.1 and 2.2 stained with phalloidin and anti-perforin to visualize the F-actin network and lytic granules, respectively. (**C**) Lysis of K562 target cells by patient and control NK cells with and without IL-2 stimulation from two independent experiments. (**D**) Lytic units required for 10% lysis of K562-target cell population, calculated from bottom panel of (C).

**Figure S5.**
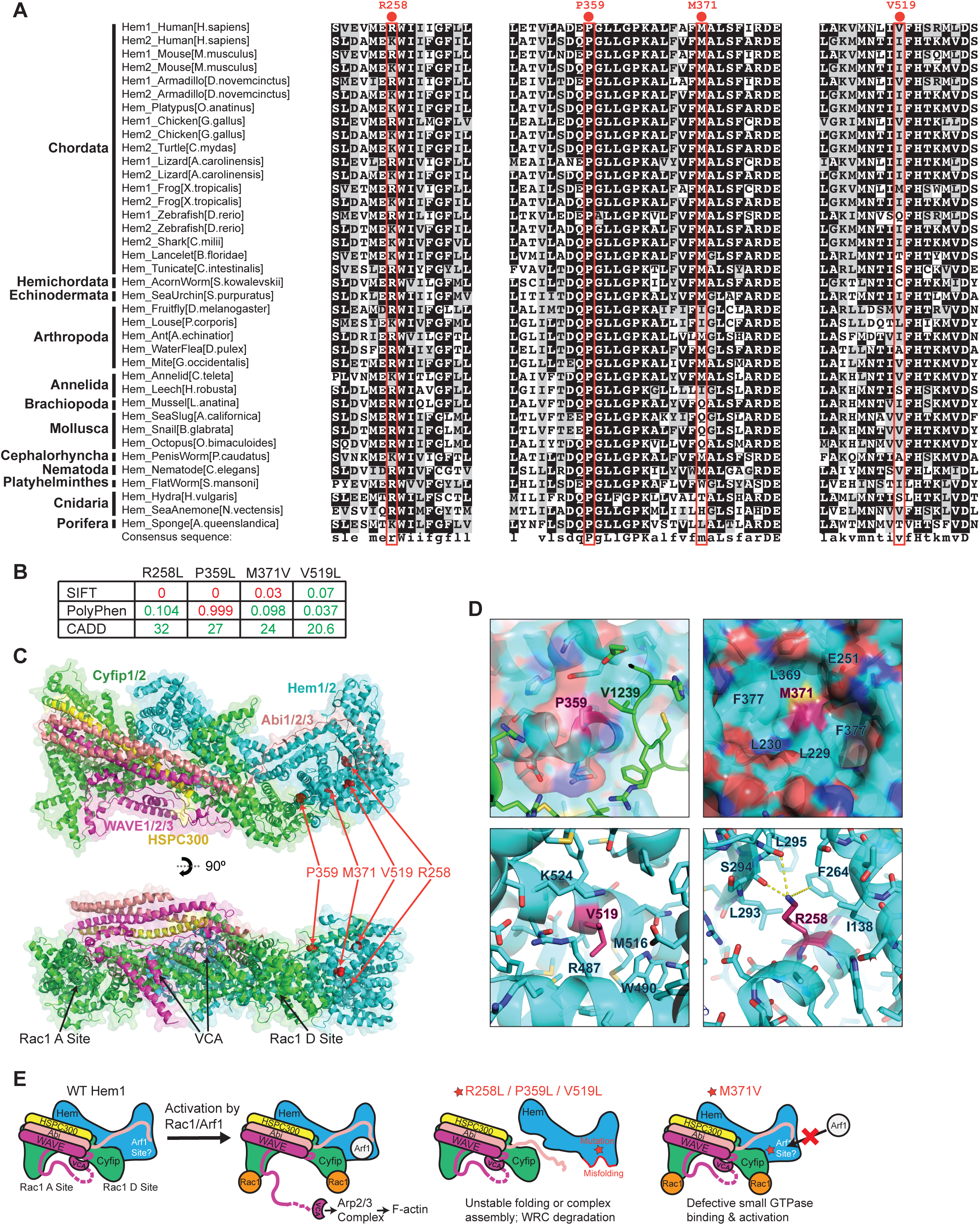
Patient NCKAP1L mutations fall within conserved regions and have detrimental effects on structure/function of the WRC. (**A**) Multiple sequence alignment of *NCKAP1L* (Hem1) and *NCKAP1* (Hem2) from various animal species, with locations of patient-derived mutations indicated in red. (**B**) Deleterious predictions of various algorithms (ref) for each of the patient mutations. Red numbers indicate that the patient mutation is above the cut-off to be considered deleterious by the indicated program. (**C**) Location of the patient mutations in Hem1 in relationship to the overall structure of the WRC. (**D**) Local structures of Hem1 patient mutations. Residues (shown in sticks) in close contact with the mutated amino acids (in purple) are indicated. (**E**) Schematic showing the impacts of different Hem1 mutations on the structure and function of the WRC.

**Figure S6.**
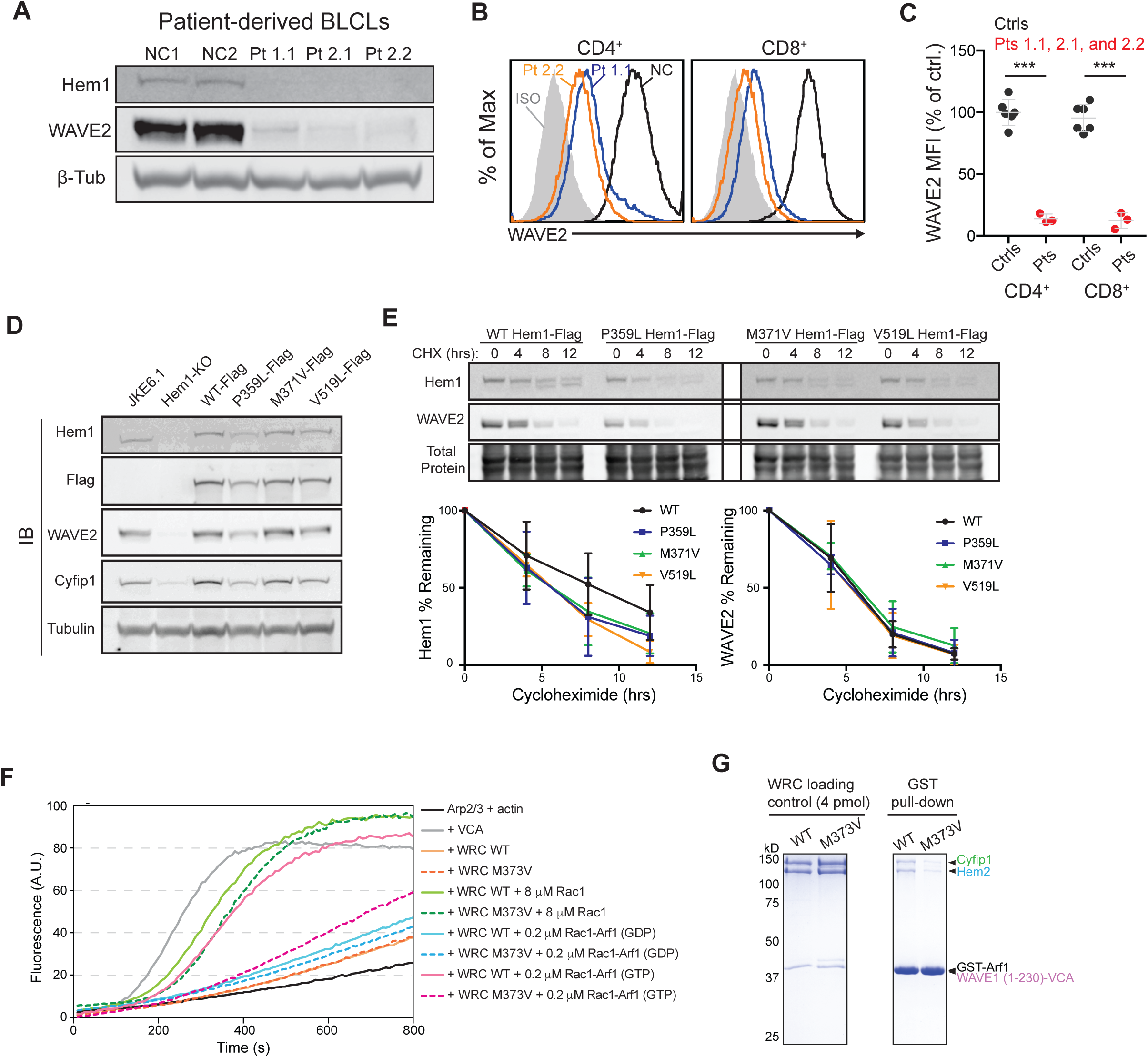
Patient mutations result disrupt assembly or function of the WRC. (**A**) Western of WRC components in lysates derived from BLCLs from patient or healthy control PBMCs. (**B**) Representative flow plots showing Wave2 expression in CD4 or CD8 T cell blasts derived from patient or normal control PBMCs. **(C)** Quantification of Wave2 MFI in patient and control CD4^+^ or CD8^+^ T cell blasts as measured by flow cytometry. (**D**) Western of lysates derived from JKE6.1 cells, a single cell JKE6.1 clone deficient for Hem1, and that clone stably reconstituted with either WT or the indicated Hem1 construct. (**E**) Representative western blot (top) and quantification (bottom) of JKE6.1 cells stably expressing the indicated Hem1-construct showing residual Hem1 and Wave2 following acute inhibition of new protein synthesis with cycloheximide. (**F**) Representative result (expansion of Fig. 2E) of pyrene-actin polymerization assays showing additional stimulations and controls comparing the WRC containing WT or M373V Hem2. Reactions contain 4 μM actin (5% pyrene labeled), 10 nM Arp2/3 complex, 100 nM WRC230VCA or VCA, and indicated amounts of Rac1 (Q61L/P29S) or Rac1 (Q61L/P29S)-Arf1 heterodimer with Arf1 pre-loaded with GDP or the unhydrolyzable GTP analog GMPPNP. (**G**) Representative Coomassie-blue stained SDS PAGE gel (expansion of Fig. 2F) showing loading controls and GST-Arf1 pull-down of the WRC230VCA containing Hem2 in the presence of active Rac1 (Q61L/P29S). \*\**P ≤* 0.01, \*\*\**P ≤* 0.001.

**Figure S7.**
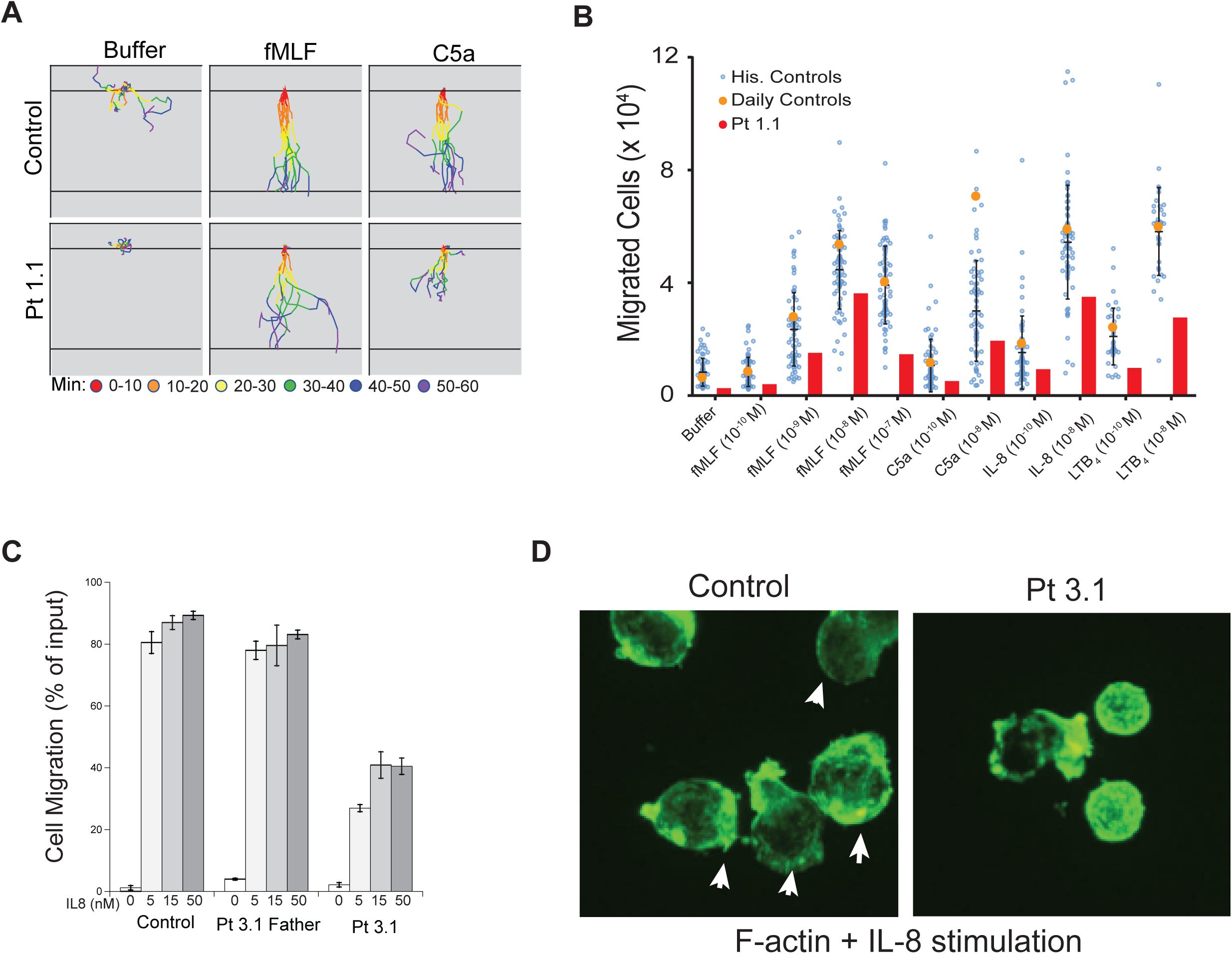
Chemotaxis is impaired in patient neutrophils. (**A**) Individual tracks of cells migrating in response to the labeled chemoattractant (expansion of Fig. 2J), color-coded to show the cellular displacement at different time intervals. (**B**) Modified Boyden chamber migration assay comparing Pt 1.1 neutrophils to daily control and historic controls for a range of chemoattractants. (**C**) Transwell migration of Pt 3.1 PBMNs in response to the indicated dose of IL-8 in comparison to a healthy donor and the patient’s M371V/WT heterozygous father. (**D**) Patient and control neutrophils stimulated with IL-8 and prepared by cytospin and stained with phalloidin to image the F-actin network. White arrows mark clear lamellipodia formation and polarization of F-actin.

**Figure S8.**
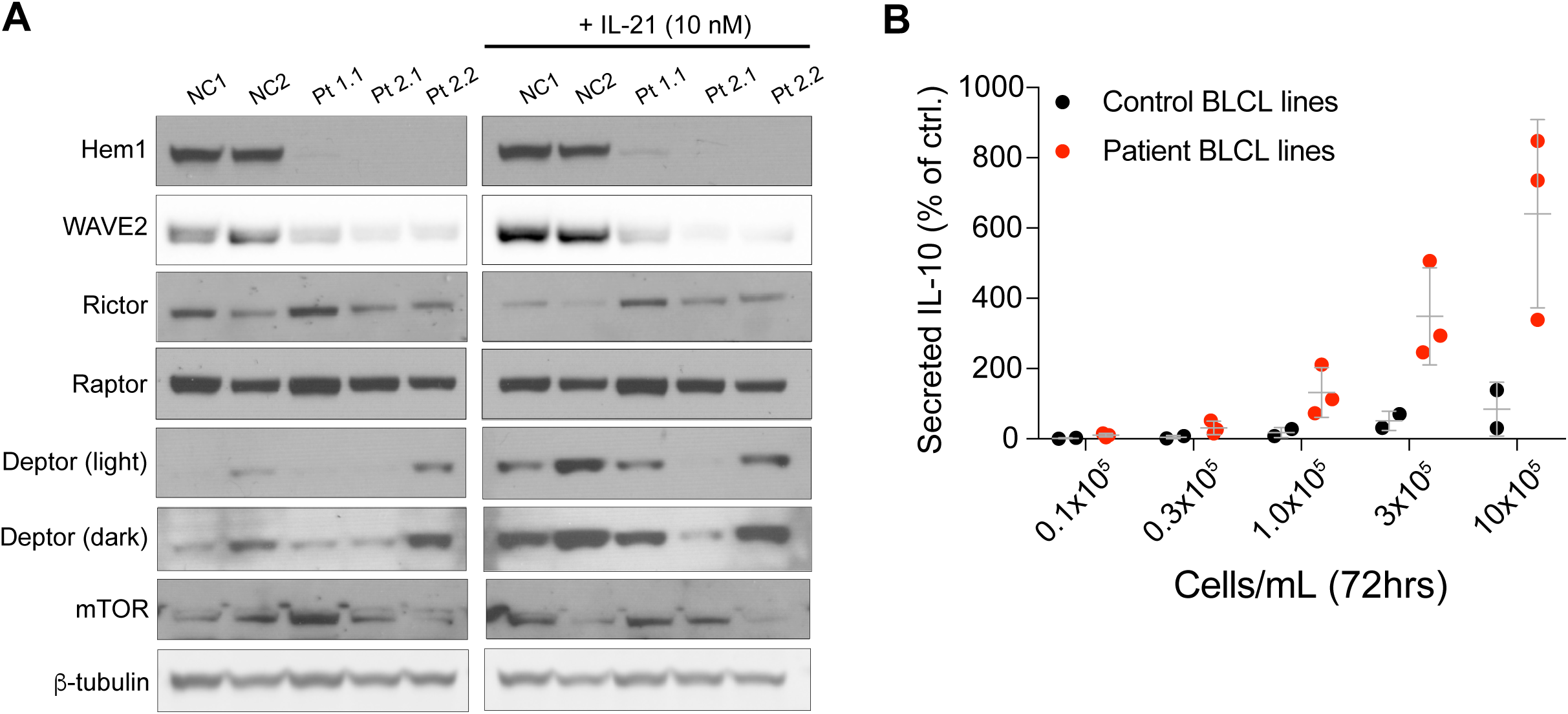
Patient BLCLs lack Hem1 and Wave2 and constitutively secrete excess cytokines. (**A**) Immunoblots of EBV-transformed BLCL lines showing WRC and mTORC2 components from 2 normal controls and 3 patients grown in either complete RPMI or RPMI supplemented with 10nM IL-21 for 72hrs prior to lysate preparation. (**B**) Measurement of IL-10 secreted by BLCL lines.

**Figure S9.**
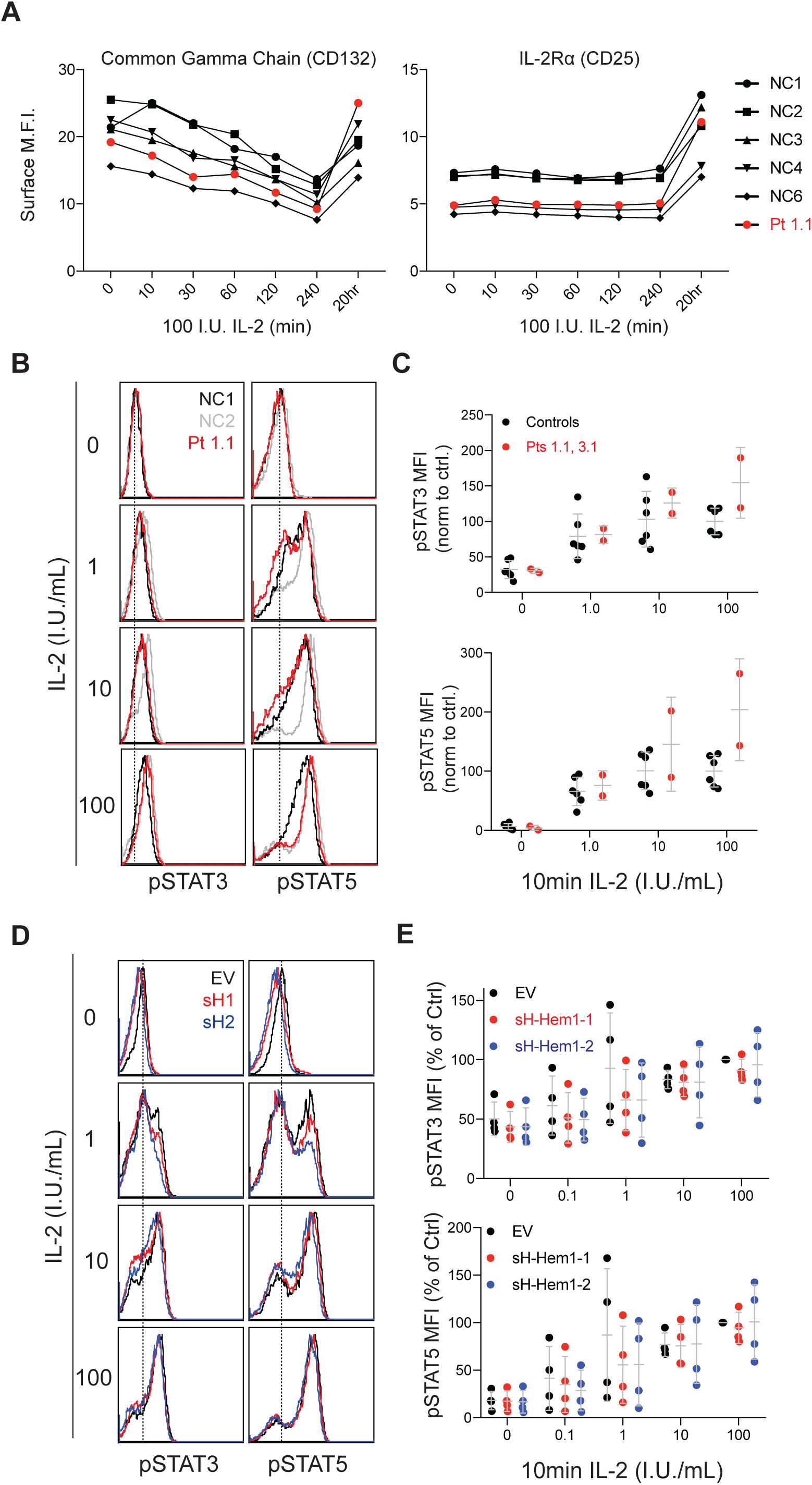
Hem1-deficiency does not affect proximal signaling downstream of the IL-2 receptor. (**A**) CD132 and CD25 surface expression following addition of 100 I.U./mL human IL-2 to patient and control cells following a 36-hour rest in complete RPMI without IL-2. (**B**) Representative flow plots and (**C**) combined results from 3 independent experiments (Pt1.1 and 3.1) showing induction of STAT3 and STAT5 phosphorylation after 10-minute stimulation with the indicated amount of IL-2. (**D**) Representative flow plots and (**E**) combined results from 4 independent experiments showing induction of STAT3 and STAT5 phosphorylation after 10minutes stimulation with the indicated amount of IL-2 in CD4 cells derived from normal donors and transduced with either shRNA against Hem1 or empty vector (EV).

**Figure S10.**
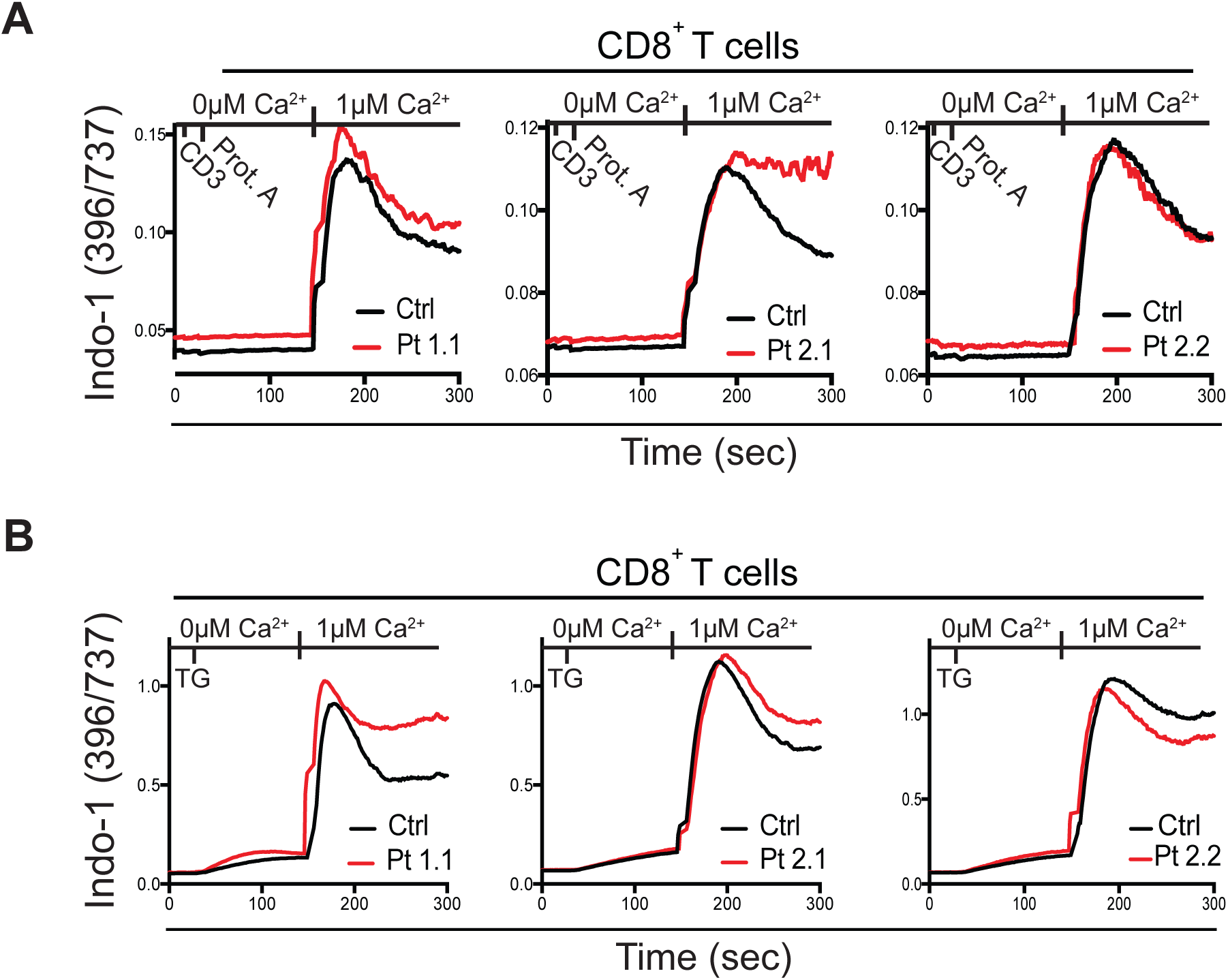
Patient cells have intact early TCR-induced signaling events. (**A**) Calcium flux in patient and control T cell blasts following stimulation with anti-CD3 and Protein A crosslinking in the presence or absence of extracellular calcium as indicated. (**B**) Calcium flux in patient and control T cell blasts following thapsigargin (TG)-induced depletion of ER-calcium stores.

**Figure S11.**
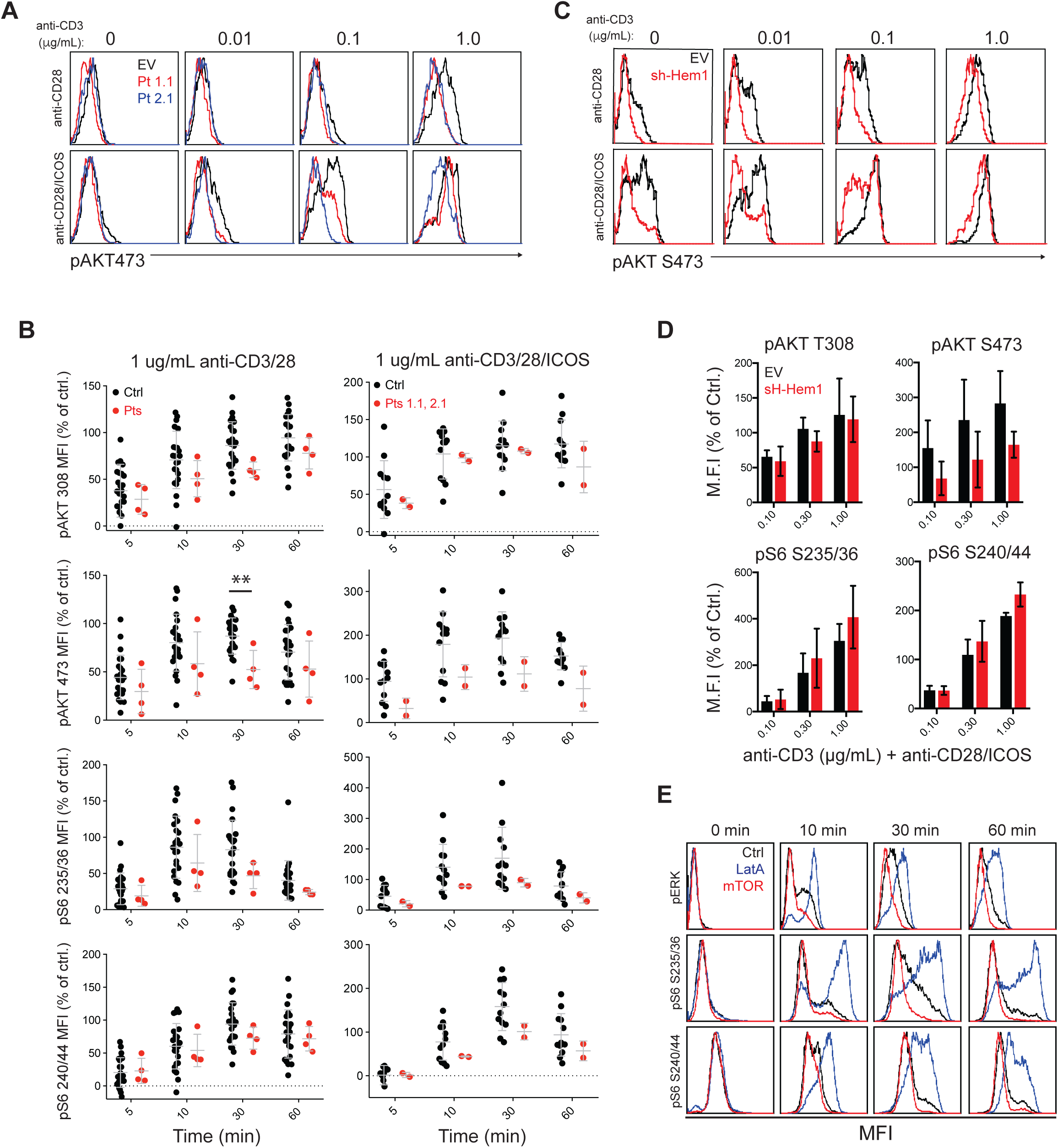
AKT and S6 phosphorylation in patient and control cells. (**A**) Extended dose response from Fig. 4C. (**B**) Time course of AKT and S6 phosphorylation in patient or normal nontrol cells stimulated with 1 ug/mL anti-CD3/CD28 with or without 1 ug/mL anti-ICOS, as measured by flow cytometry. Shown are the MFI values for each phospho-protein stain normalized to the maximal control response. (**C**) Extended dose response from Fig. 4C. (**D**) Average and SD of multiple experiments stimulated as in Fig. S11C and stained for the indicated phosphor-protein. (**E**) Representative flow plots of ERK and S6 phosphorylation following stimulation of normal control T cell blasts as in (extension of Fig. 4E).

**Figure S12.**
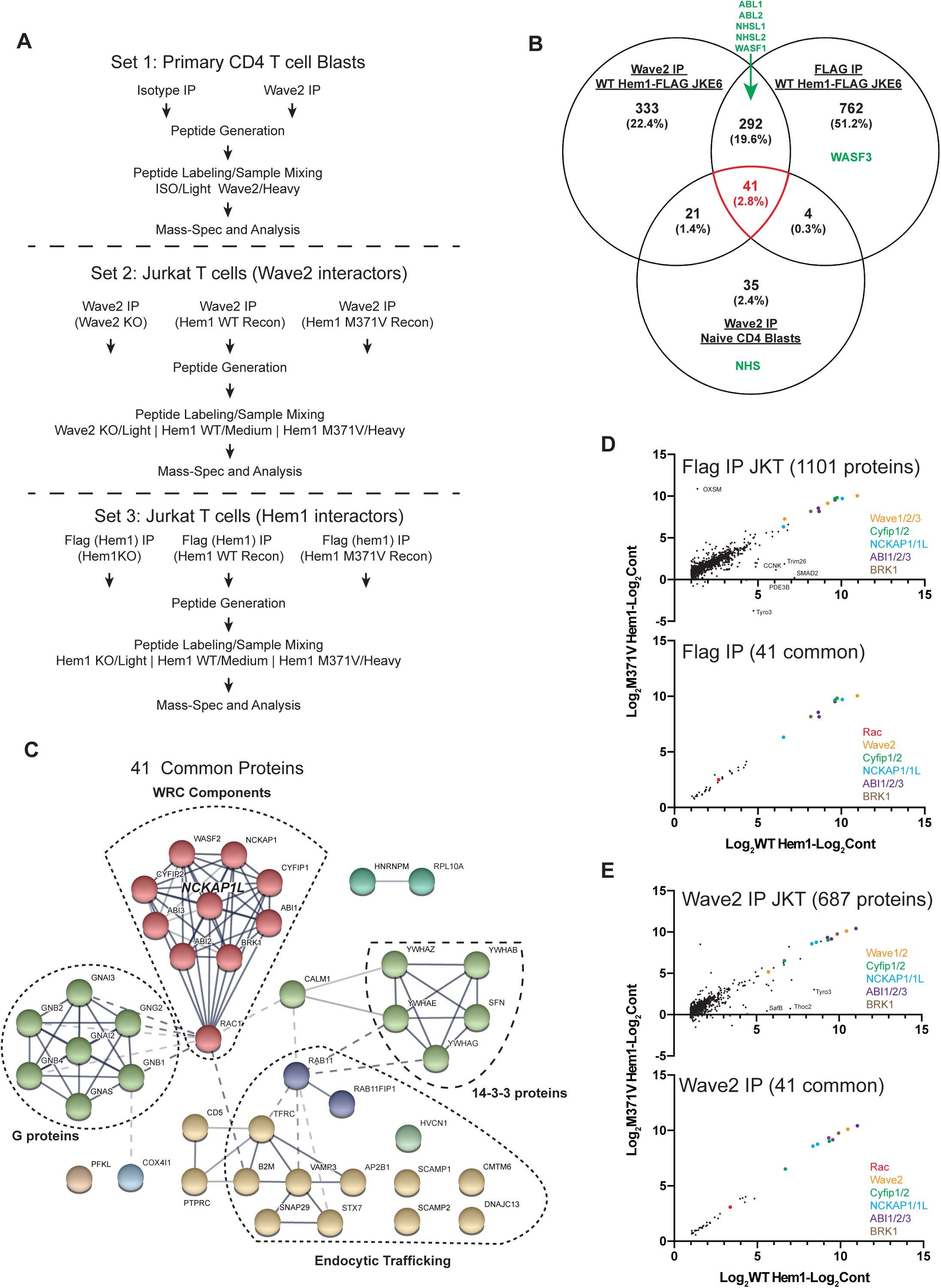
IP-Mass spec analysis of Wave2, Hem1-WT, and Hem1-M371V interacting proteins. (**A**) Work flow of IP-Mass Spectrometry experiments. (**B**) Venn Diagram showing number of commonly identified and unique proteins. Proteins that are known to interact with the WRC but were not identified in all three experimental setups are shown in green. (**C**) STRING Protein-Protein Interaction mapping on the 41 proteins identified in all three immunoprecipitations. (**D**) Relative abundance of proteins identified by quantitative mass-spec in the Flag IP in Jurkat cells from the WT and M371V Hem1 variants (top) or limiting the comparison to those proteins identified in all three IP-mass spec experiments (bottom). (**E**) Relative abundance of proteins identified by quantitative mass-spec in the Wave2 IP in Jurkat cells from the WT and M371V Hem1 variants (top) or limiting the comparison to those proteins identified in all three IP-mass spec experiments (bottom).

**Figure S13.**
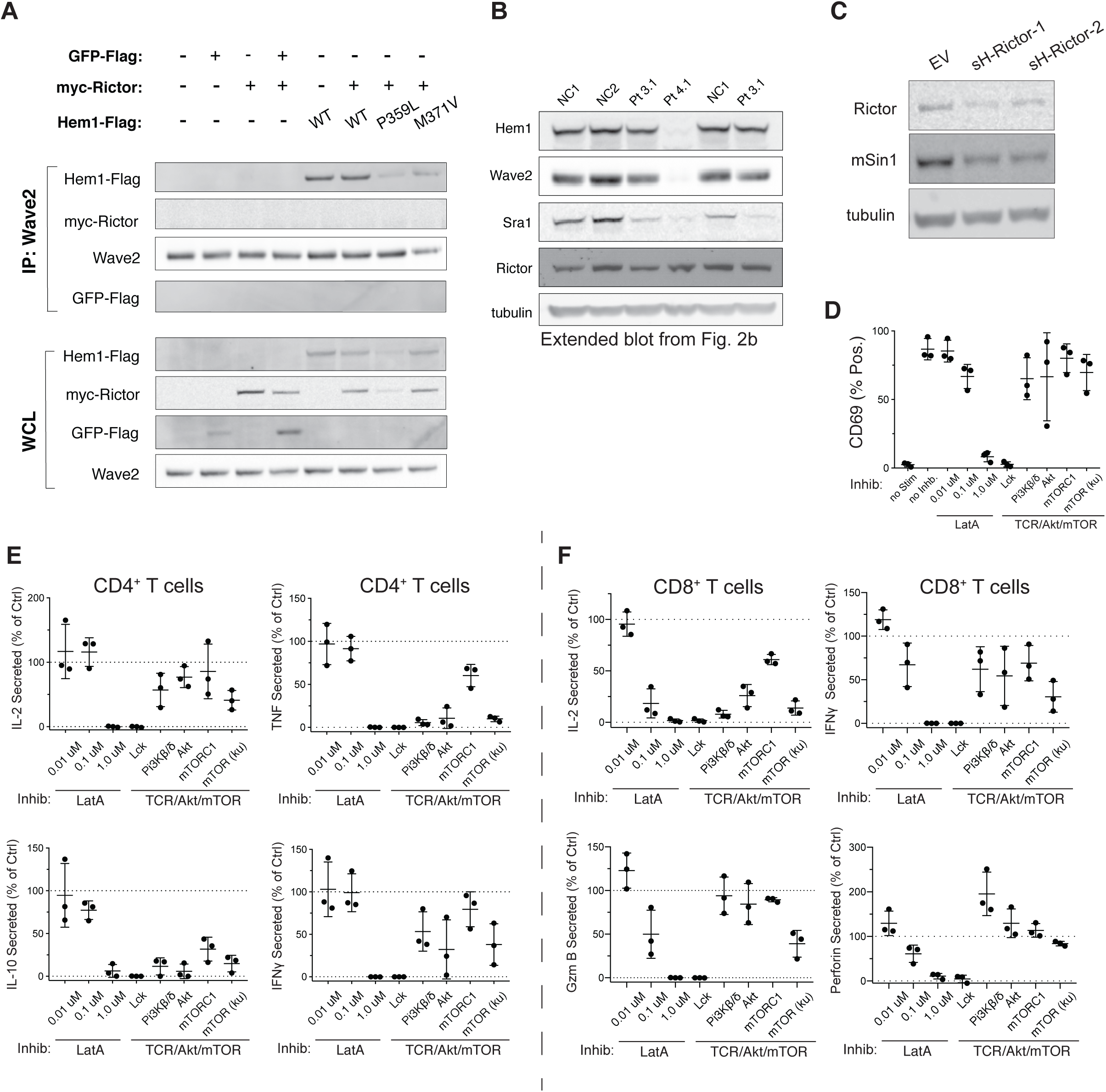
Rictor is expressed in patient T cells and does not interact with Wave2 and PI3K/mTORC2/AKT inhibitors selectively alter cytokine production. (**A**) Immunoprecipitation of Wave2 followed by immunoblot for interacting proteins. (**B**) WRC component and Rictor expression in patient and normal control cells. Blot is extended from Fig. 2B. (**C**) Immunoblot showing efficiency of shRNA-mediated Rictor knockdown in naïve CD4^+^ T cells relative to empty vector (EV) control. (**D**) CD69 expression on naïve CD4^+^ T cells from a normal donor stimulated with plate-bound anti-CD3/28/ICAM1 (1 ug/mL each) in the presence of the indicated inhibitors for 36 hours. (**E**) Cytokine production in CD4^+^ cells following an 18-hour stimulation with anti-CD3/28/ICAM1 (1 ug/ml each) coated surfaces and co-treatment with the indicated inhibitors. (**F**) Cytokine production in CD8^+^ cells following treatment as in (E).

**Table S1. Clinical and functional characteristics of patients with Hem1-LOF mutations.**

N/D indicates test was not done.

**Table S2. Evaluation of key immunological subsets**

**Table S3. Homozygous variants tracking with patient disease**

A list of homozygous variants of unknown significance identified by WES in the patient population.

**Table S4. IP/Mass-spec analysis of Hem1 and Wave2 interacting proteins**

**(Tab1)** Proteins identified from Wave2 IP or Flag IP from either JKE6.1 cells knocked out for Wave2 or Wave2, Hem1-KO JKE6.1 cells reconstituted with WT Hem1-Flag or M371V Hem1-Flag, digested, and then labeled with Low, Med, and High isotope tags, respectively. **(Tab2)** Proteins identified from Isotype or Wave2 IP from primary CD4^+^ T cell blasts derived from PBMCs of a healthy donor. **(Tab3)** Proteins identified from Wave2 IP from either JKE6.1 cells knocked out for Wave2, Hem1-KO JKE6.1 cells reconstituted with WT Hem1-Flag or M371V Hem1-Flag, digested, and then labeled with Low, Med, and High tags, respectively. Proteins above a Ratio of 1.5 and Log_2_ difference of 1.0 are shown. **(Tab4)** Proteins identified from Flag IP from either JKE6.1 cells knocked out for Hem1, Hem1-KO JKE6.1ells reconstituted with WT Hem1-Flag or M371V Hem1-Flag, digested, and then labeled with Low, Med, and High epitope tags, respectively. Proteins above a Ratio of 1.5 and Log_2_ difference of 1.0 are shown. **(Tab5)** Proteins identified from Isotype or Wave2 IP from primary CD4^+^ T cell blasts derived from PBMCs of a healthy donor. Proteins above a ratio of 1.5 and a probability score of 1.0 are shown. **(Tab6)** A list of proteins positively identified as interacting with both Hem1 and Wave2 in Jurkat Cells and Wave2 in primary cells.

**Movie S1.**

Primary CD4^+^ T cell blasts from a normal control or Pt 1.1 transfected with Lifeact-GFP and spreading on stimulatory coverslips coated with 1 ug/mL antiCD3/28 and 1 ug/mL ICAM-1.

**Movie S2.**

Hem-1 deficient clone derived from JKE6.1 parental cells, and transduced with Lifeact-GFP expressing virus as well as either WT Hem1-Flag, P359VLHem1-Flag, M371V Hem1-Flag, or V519L Hem1-Flag. Cells are spreading on stimulatory coverglass coated with 1 ug/mL antiCD3 and 1 ug/mL ICAM-1

**Movie S3.**

Normal control or patient CD4^+^ T cells migrating spontaneously on coverslips coated with 1 ug/mL ICAM-1.

**Movie S4.**

Normal control neutrophils (top) or patient 1.1 neutrophils (bottom) migrating in an EZ-TAXIScan chemotaxis chamber in response to either no chemoattractant (left), fMLF (center), or C5a (right).

## Work Cited

1. C. Cunningham-Rundles, P. P. Ponda, Molecular defects in T- and B-cell primary immunodeficiency diseases. Nat Rev Immunol 5, 880–892 (2005).

2. H. K. Lehman, Autoimmunity and Immune Dysregulation in Primary Immune Deficiency Disorders. Curr Allergy Asthma Rep 15, 53 (2015).

3. B. Chen, S. B. Padrick, L. Henry, M. K. Rosen, Biochemical reconstitution of the WAVE regulatory complex. Methods Enzymol 540, 55–72 (2014).

4. T. E. Stradal et al., Regulation of actin dynamics by WASP and WAVE family proteins. Trends Cell Biol 14, 303–311 (2004).

5. J. C. Nolz et al., The WAVE2 complex regulates T cell receptor signaling to integrins via Abl- and CrkL-C3G-mediated activation of Rap1. J Cell Biol 182, 1231–1244 (2008).

6. L. Shao et al., The Wave2 scaffold Hem-1 is required for transition of fetal liver hematopoiesis to bone marrow. Nat Commun 9, 2377 (2018).

7. B. Chen et al., Rac1 GTPase activates the WAVE regulatory complex through two distinct binding sites. Elife 6, (2017).

8. B. Chen et al., The WAVE regulatory complex links diverse receptors to the actin cytoskeleton. Cell 156, 195–207 (2014).

9. V. Koronakis et al., WAVE regulatory complex activation by cooperating GTPases Arf and Rac1. Proc Natl Acad Sci U S A 108, 14449–14454 (2011).

10. Z. Chen et al., Structure and control of the actin regulatory WAVE complex. Nature 468, 533–538 (2010).

11. A. M. Lebensohn, M. W. Kirschner, Activation of the WAVE complex by coincident signals controls actin assembly. Mol Cell 36, 512–524 (2009).

12. Y. Kim et al., Phosphorylation of WAVE1 regulates actin polymerization and dendritic spine morphology. Nature 442, 814–817 (2006).

13. R. A. Saxton, D. M. Sabatini, mTOR Signaling in Growth, Metabolism, and Disease. Cell 169, 361–371 (2017).

14. D. A. Guertin et al., Ablation in mice of the mTORC components raptor, rictor, or mLST8 reveals that mTORC2 is required for signaling to Akt-FOXO and PKCalpha, but not S6K1. Dev Cell 11, 859–871 (2006).

15. Z. Zou et al., mTORC2 promotes cell survival through c-Myc-dependent up-regulation of E2F1. J Cell Biol 211, 105–122 (2015).

16. K. Lee et al., Mammalian target of rapamycin protein complex 2 regulates differentiation of Th1 and Th2 cell subsets via distinct signaling pathways. Immunity 32, 743–753 (2010).

17. L. A. Van de Velde, P. J. Murray, Proliferating Helper T Cells Require Rictor/mTORC2 Complex to Integrate Signals from Limiting Environmental Amino Acids. J Biol Chem 291, 25815–25822 (2016).

18. M. Lek et al., Analysis of protein-coding genetic variation in 60,706 humans. Nature 536, 285–291 (2016).

19. H. Park et al., A point mutation in the murine Hem1 gene reveals an essential role for Hematopoietic protein 1 in lymphopoiesis and innate immunity. J Exp Med 205, 2899–2913 (2008).

20. J. C. Nolz et al., The WAVE2 complex regulates actin cytoskeletal reorganization and CRAC-mediated calcium entry during T cell activation. Curr Biol 16, 24–34 (2006).

21. O. D. Weiner et al., Hem-1 complexes are essential for Rac activation, actin polymerization, and myosin regulation during neutrophil chemotaxis. PLoS Biol 4, e38 (2006).

22. S. Murugesan et al., Formin-generated actomyosin arcs propel T cell receptor microcluster movement at the immune synapse. J Cell Biol 215, 383–399 (2016).

23. E. Derivery et al., Free Brick1 is a trimeric precursor in the assembly of a functional wave complex. PLoS One 3, e2462 (2008).

24. J. S. Orange et al., Wiskott-Aldrich syndrome protein is required for NK cell cytotoxicity and colocalizes with actin to NK cell-activating immunologic synapses. Proc Natl Acad Sci U S A 99, 11351–11356 (2002).

25. F. Tamzalit et al., Interfacial actin protrusions mechanically enhance killing by cytotoxic T cells. Sci Immunol 4, (2019).

26. A. Leithner et al., Diversified actin protrusions promote environmental exploration but are dispensable for locomotion of leukocytes. Nat Cell Biol 18, 1253–1259 (2016).

27. C. Basquin et al., Membrane protrusion powers clathrin-independent endocytosis of interleukin-2 receptor. EMBO J 34, 2147–2161 (2015).

28. A. T. Ritter et al., Cortical actin recovery at the immunological synapse leads to termination of lytic granule secretion in cytotoxic T lymphocytes. Proc Natl Acad Sci U S A 114, E6585–E6594 (2017).

29. D. D. Sarbassov, D. A. Guertin, S. M. Ali, D. M. Sabatini, Phosphorylation and regulation of Akt/PKB by the rictor-mTOR complex. Science 307, 1098–1101 (2005).

30. L. R. Pearce et al., Identification of Protor as a novel Rictor-binding component of mTOR complex-2. Biochem J 405, 513–522 (2007).

31. A. Diz-Munoz et al., Membrane Tension Acts Through PLD2 and mTORC2 to Limit Actin Network Assembly During Neutrophil Migration. PLoS Biol 14, e1002474 (2016).

## References

7. Z. Chen et al., Structure and control of the actin regulatory WAVE complex. Nature 468, 533–538 (2010).

8. B. Chen et al., Rac1 GTPase activates the WAVE regulatory complex through two distinct binding sites. Elife 6, (2017).

9. B. Chen et al., The WAVE regulatory complex links diverse receptors to the actin cytoskeleton. Cell 156, 195–207 (2014).

10. V. Koronakis et al., WAVE regulatory complex activation by cooperating GTPases Arf and Rac1. Proc Natl Acad Sci U S A 108, 14449–14454 (2011).

32. K. B. Sanborn et al., Myosin IIA associates with NK cell lytic granules to enable their interaction with F-actin and function at the immunological synapse. J Immunol 182, 6969–6984 (2009).

33. K. B. Sanborn, G. D. Rak, A. N. Mentlik, P. P. Banerjee, J. S. Orange, Analysis of the NK cell immunological synapse. Methods Mol Biol 612, 127–148 (2010).

34. P. P. Banerjee, J. S. Orange, Quantitative measurement of F-actin accumulation at the NK cell immunological synapse. J Immunol Methods 355, 1–13 (2010).

35. J. Schindelin et al., Fiji: an open-source platform for biological-image analysis. Nat Methods 9, 676–682 (2012).

36. B. Chen, S. B. Padrick, L. Henry, M. K. Rosen, Biochemical reconstitution of the WAVE regulatory complex. Methods Enzymol 540, 55–72 (2014).

37. B. Chen et al., Rac1 GTPase activates the WAVE regulatory complex through two distinct binding sites. Elife 6, (2017).

38. E. K. O’Shea, K. J. Lumb, P. S. Kim, Peptide ‘Velcro’: design of a heterodimeric coiled coil. Curr Biol 3, 658–667 (1993).

39. E. C. Keilhauer, M. Y. Hein, M. Mann, Accurate protein complex retrieval by affinity enrichment mass spectrometry (AE-MS) rather than affinity purification mass spectrometry (AP-MS). Mol Cell Proteomics 14, 120–135 (2015).

40. P. J. Boersema, R. Raijmakers, S. Lemeer, S. Mohammed, A. J. Heck, Multiplex peptide stable isotope dimethyl labeling for quantitative proteomics. Nat Protoc 4, 484–494 (2009).

41. J. Cox, M. Mann, MaxQuant enables high peptide identification rates, individualized p.p.b.-range mass accuracies and proteome-wide protein quantification. Nat Biotechnol 26, 1367–1372 (2008).

42. Y. Lin et al., Sodium laurate, a novel protease- and mass spectrometry-compatible detergent for mass spectrometry-based membrane proteomics. PLoS One 8, e59779 (2013).

43. J. Rappsilber, M. Mann, Y. Ishihama, Protocol for micro-purification, enrichment, pre-fractionation and storage of peptides for proteomics using StageTips. Nat Protoc 2, 1896–1906 (2007).

